# Conformational activation of GSK3β by an environmental toxicant suppresses hedgehog signalling and disrupts ciliary homeostasis

**DOI:** 10.64898/2026.08.03.742442

**Authors:** Ravikant Piyush, Himani Barmola, Anwesha Bhattacharjya, Abhinn Gupta, Prasenjit Bhaumik, Sathees C Raghavan, Bibha Choudhary, Sudarshan Gadadhar, Satyanarayan Rao, Swapnil Rohidas Shinde

## Abstract

Primary cilium-dependent hedgehog signalling is essential for embryonic development, tissue patterning, and organ homeostasis, and its dysfunction causes ciliopathies, a clinically diverse spectrum of developmental and reproductive disorders. Whether environmental chemicals can phenocopy genetic ciliopathies by directly targeting ciliary kinase machinery has remained unknown. Here we show that endosulfan, a banned organochlorine pesticide linked to congenital and reproductive defects, suppresses hedgehog signalling by driving proteolytic processing of GLI transcription factors into repressor forms. Excluding ciliary receptor trafficking, cAMP signalling, and GLI–DNA binding, we identify PKA and GSK3β as direct endosulfan targets: endosulfan allosterically fine-tunes PKA activity and, to our knowledge, is the first reported small-molecule activator of GSK3β, stabilising its active conformation, a profile distinct from all known inhibitors. We further identify *Cetn3* and *Cep250* as novel GLI-regulated genes required for centriole cohesion, both of which are repressed upon endosulfan exposure, linking this kinase axis to the reproductive defects reported in exposed human populations and animal models. These findings establish a chemical–biological axis through which an environmental toxicant hijacks core kinase signalling to phenocopy a genetic ciliopathy.

## Introduction

Hedgehog (HH) signalling governs embryonic patterning, tissue morphogenesis, and adult stem cell homeostasis across vertebrates, and its disruption can produce a clinically diverse spectrum of developmental disorders, including neural tube defects, skeletal anomalies, reproductive failure, and congenital malformations [1,2]. In vertebrates, hedgehog signalling is transduced through primary cilia, evolutionarily conserved, microtubule-based organelles that project from the apical surface of most post-mitotic cells and serve as essential signalling hubs [3]. Primary cilia host the HH receptor Patched 1 (PTCH1) and the G protein-coupled receptors Smoothened (SMO) and GPR161, whose coordinated redistribution controls the processing and nuclear translocation of GLI transcription factors [4]. In the absence of HH ligands, cyclic AMP-dependent protein kinase A (PKA), glycogen synthase kinase 3β (GSK3β), and casein kinase 1 (CK1) sequentially phosphorylate GLI2 and GLI3, driving their partial proteasomal degradation into transcriptional repressors (GLI^R^) that silence pathway target genes [4–6]. HH ligand binding blocks this phosphorylation cascade, enabling full-length GLI (GLIFL) accumulation, nuclear translocation, and target gene activation [7–9]. Despite the central importance of PKA and GSK3β in regulating GLI processing, the mechanisms by which their activities are modulated in the context of hedgehog signalling and whether environmental chemicals can perturb this kinase machinery to disrupt pathway output remain incompletely understood.

GSK3β is a constitutively active, dual-specificity kinase whose activity is regulated primarily through inhibitory phosphorylation at Ser9 and activating autophosphorylation at Tyr216 [10]. It occupies a central position in multiple signalling networks beyond the Hedgehog pathway, including Wnt, insulin, and neurotrophin pathways, and its dysregulation is implicated in Alzheimer’s disease, type 2 diabetes, bipolar disorder, and cancer [11]. Extensive therapeutic effort has focused on GSK3β inhibition; selective inhibitors, including lithium, SB-216763, and BRD3731, are widely used as research tools, and some have entered clinical evaluation. By contrast, no small-molecule activator of GSK3β has been reported to date, in part because the constitutively active nature of the kinase has historically made activation seem pharmacologically unnecessary. Whether the active conformation of GSK3β can be further stabilised by small-molecule binding, and what the consequences of such stabilisation would be for hedgehog pathway regulation and downstream developmental gene expression, remain open questions.

The possibility that environmental chemicals could disrupt the hedgehog pathway by targeting its kinase machinery was raised by the discovery of cyclopamine. This plant-derived alkaloid causes teratogenic cyclopia in livestock by directly inhibiting SMO [12,13]. This precedent, an environmental small molecule that produces developmental defects through specific antagonism of the hedgehog pathway, raises the question of whether unrelated environmental chemicals might similarly perturb pathway activity through distinct molecular mechanisms. Endosulfan (ES), an affordable broad-spectrum organochlorine insecticide, provides a compelling case study [14]. Despite its 2011 ban under the Stockholm Convention [15], endosulfan continues to be used in some developing countries, and its metabolites bioaccumulate in human blood, adipose tissue, placenta, and breast milk [16–22]. The Kerala tragedy illustrates the human health consequences of sustained exposure. According to the 2019 Kerala state government report, over 6,700 individuals suffer from endosulfan-related conditions and more than 770 have died from poisoning. Epidemiological studies from 2012 and 2019 document a striking phenotypic spectrum in exposed individuals: congenital malformations, cerebral palsy, skeletal anomalies, reproductive failure, and skin disorders that overlap substantially with the phenotypes produced by genetic disruption of Hedgehog signalling and primary cilia function [23–25]. In exposed males, altered sperm morphology and reduced sperm counts predominate; in females, menstrual irregularities, foetal death, and gynaecological complications are most common [26,27]. Comparable reproductive and developmental phenotypes are recapitulated in endosulfan-exposed mice and zebrafish [27,28].

Most clinical manifestations of endosulfan exposure have been attributed to its well-documented neurotoxicity through acetylcholinesterase inhibition. However, the breadth and developmental specificity of the phenotypic spectrum, particularly the congenital malformations and reproductive defects, prompted us to hypothesise that endosulfan may additionally perturb primary cilia-mediated hedgehog signalling, whose disruption produces a remarkably overlapping set of phenotypes in both human genetic disease and animal models **(Table EV1 and EV2)**.

Here we show that endosulfan suppresses hedgehog signalling across multiple ciliated cell types and in the gonads of exposed mice by driving GLI3 processing into its repressor form and suppressing GLI1 expression. Through systematic exclusion of upstream pathway components, we identify PKA and GSK3β as direct endosulfan targets. Endosulfan allosterically fine-tunes PKA activity and, in a pharmacologically unprecedented finding, acts as the first reported small-molecule activator of GSK3β, stabilising the active kinase conformation through a conformational selection mechanism distinct from all known GSK3β inhibitors. Critically, endosulfan does not activate the structurally related kinase CDK1 and does not form stable complexes with CDK1 or CDK2, establishing kinase selectivity. Beyond canonical hedgehog targets, we identify *Cetn3* and *Cep250* as novel GLI-regulated genes that govern centriole cohesion, both of which are repressed by endosulfan, providing a candidate molecular explanation for the reproductive phenotypes associated with exposure.

## Results

### Endosulfan disrupts primary cilia homeostasis and GLI processing

Primary cilia are essential for hedgehog signalling in vertebrates as they host and regulate key pathway components. To test whether endosulfan disrupts primary cilia-hedgehog homeostasis, we first established non-cytotoxic treatment conditions across the cell lines used in this study. We determined the IC_50_ of endosulfan in IMCD3 cells (33.82 µM) and used a concentration lower than half of this throughout the study (**Figure EV1A**). At this concentration, endosulfan did not affect ciliary number or cell cycle progression (**EV1B-D**). To investigate the effect of endosulfan on primary cilia, we next examined cilia length using acetylated tubulin (α-tubulin Lys40 acetylation, acTub) and ARL13B as ciliary markers. We observed that ciliary length was significantly reduced in endosulfan-treated cells compared with control cells (**Figure 1A-B**). Intraflagellar transport (IFT) machinery maintains cilia; hence, to identify the cause of this decrease in ciliary length, we next assessed ciliary IFT88 levels in IMCD3 cells and observed their significant downregulation upon endosulfan treatment (**Figure 1C-D**). We further examined the number of cilia, ciliary length, and IFT88 levels in a parallel paradigm in which cells were treated with endosulfan after cilia had formed. Using half the IC_50_ concentration (13.35 µM) determined for this paradigm (**Figure EV2A**), cilia number remained unchanged, whereas ciliary length was significantly reduced, and ciliary IFT88 levels were downregulated (**Figure EV2B-F**). We next performed the same experiments in NIH3T3 cells to assess whether the endosulfan effects on ciliation were conserved. At half the IC_50_ concentration (52.41 µM) (**Figure EV3A**), cilia number remained unchanged, whereas ciliary length was significantly reduced (**Figure EV3B-D**). Consistent with this, ciliary IFT88 levels were significantly downregulated (**Figure EV3E**), corroborating a robust effect of endosulfan on ciliary length homeostasis across cell lines.

**Figure 1.**
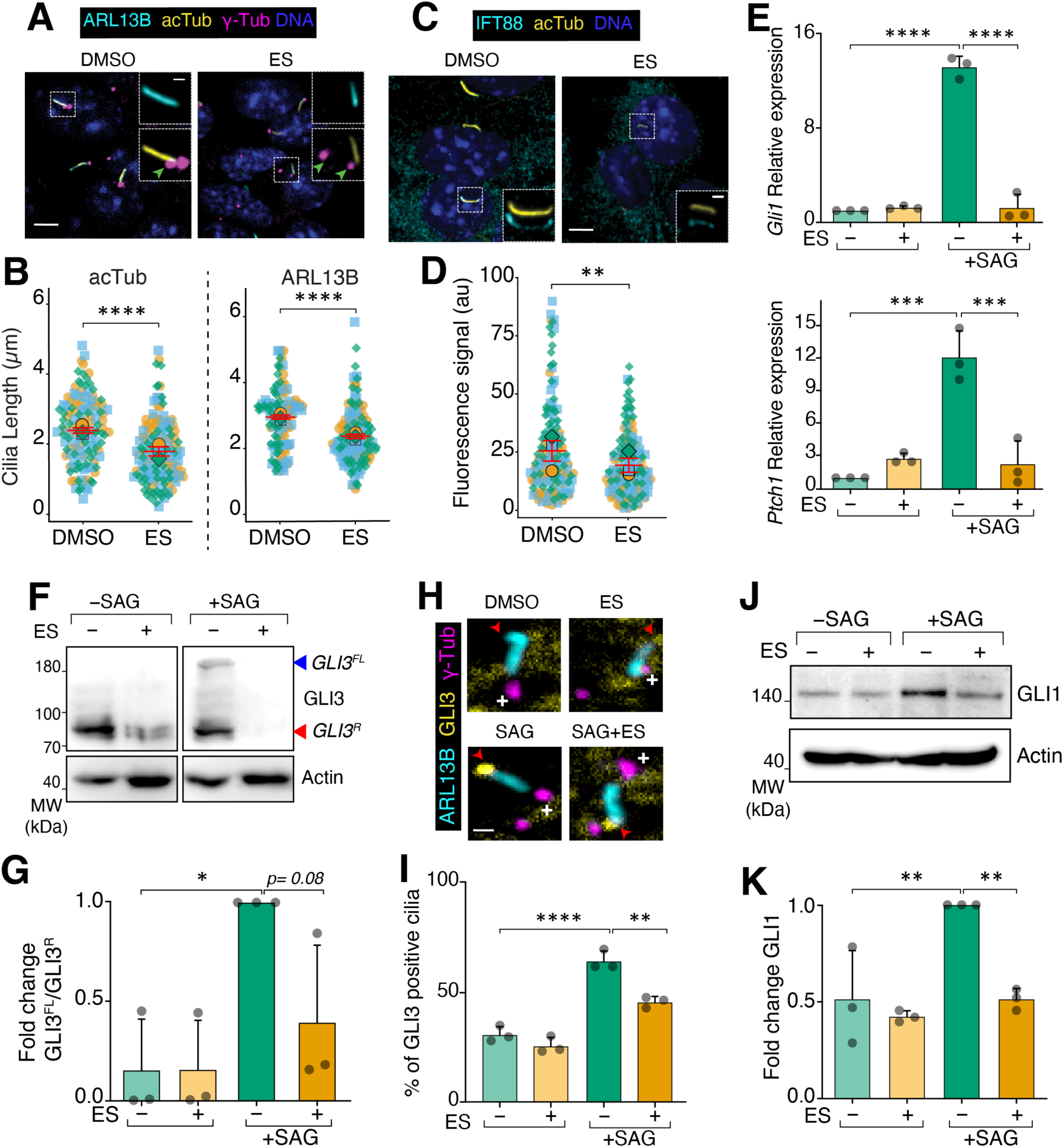
Endosulfan perturbs cilia and suppresses Hedgehog pathway activity. **(A)** IMCD3 cells were treated with DMSO or endosulfan for 48 h, then fixed and stained for ARL13B (cyan), acetylated tubulin (acTub, yellow), γ-Tubulin (magenta), and DNA (blue). Insets show a representative cilium, with the base marked by a green arrowhead. Scale bars, 5 µm (main panel), 1 µm (inset). **(B)** Quantification of ciliary length for the cilia shown in **(A)**. Ciliary length was measured using acTub; (DMSO n = 132, endosulfan n = 146) and ARL13B (DMSO n = 122, endosulfan n = 146) as independent ciliary markers. In the superplots, three independent experimental data points are represented by three different shapes, with colours indicated. Each square, circle, or triangle represents a data point, and the larger version of its shape represents the mean. Data represent all data points from three independent experiments, mean ±SEM. **(C)** DMSO- or endosulfan-treated IMCD3 cells were stained for IFT88 (cyan), acTub (yellow), and DNA (blue). **(D)** The fluorescence intensities of IFT88 are plotted in the superplots, using the same conventions as in **(B)**; (DMSO, n = 157; endosulfan, n = 157). **(E)** qRT-PCR analysis of hedgehog pathway genes in IMCD3 cells treated with SAG, endosulfan, or vehicle (0.05% DMSO). Gene expression in the DMSO-treated condition was set to 1. Data represent the mean ±SD of three independent experiments. **(F)** IMCD3 cells treated with the same paradigm as in **E** were lysed, and the lysates were probed for GLI3 and actin (loading control) via immunoblotting. **(G)** Quantification of GLI3 full-length to GLI3 repressor ratio (GLI3^FL^/GLI3^R^). Signal intensities for GLI3^FL^ or GLI3^R^ were measured and normalised against the loading control. The GLI3^FL^/GLI3^R^ ratio in the SAG-treated condition was set to 1. The experiment shown in **F** was repeated three times, and the data represent the mean ±SD of three independent experiments. **(H)** IMCD3 cells were treated as in **E**, fixed and stained for ARL13B (cyan), GLI3 (yellow), and γ-Tubulin (magenta). Representative cilia are shown for each condition; the base of the cilia is marked with γ-Tubulin (shown with +), and the tip is marked with a red arrowhead; corresponding full cell images are shown in **Figure EV1E**. Scale bar, 1 µm. **(I)** Cilia positive for GLI3 at the tip were quantified, and the percentage was plotted; data represent the mean ± SD of three independent experiments. **(J)** IMCD3 cells treated with the same paradigm as in **E** were lysed, and the lysates were probed for GLI1 and Actin (loading control) via immunoblotting. **(K)** Quantification of GLI1 levels. GLI1 signal intensities were measured and normalised against the loading control. GLI1 expression in the SAG-treated condition was set to 1. The experiment shown in **J** was repeated three times, and the data represent the mean ±SD of three independent experiments. For all the experiments, fluorescence intensities are expressed in arbitrary units (a.u.). Asterisks denote statistical significance as determined by unpaired Student’s t-test or one-way ANOVA followed by Tukey’s post-hoc multiple comparison test, as appropriate: **** p ≤ 0.0001; *** p ≤ 0.001; ** p ≤ 0.01; * p ≤ 0.05.

Having established that endosulfan reduces ciliary IFT88 levels and ciliary length, we next investigated its effect on hedgehog signalling, given the essential role of primary cilia in pathway transduction. We first examined the mRNA expression of the hedgehog target genes *Gli1* and *Ptch1* in IMCD3 cells, a well-validated cell line for primary cilia-dependent hedgehog studies. We activated hedgehog signalling by treating cells with SAG, a Smoothened agonist, for 16 hours. qRT-PCR data showed that *Gli1* and *Ptch1* expression increased significantly upon SAG treatment compared to vehicle control (**Figure 1E**); however, simultaneous treatment with SAG and endosulfan significantly decreased the expression of these hedgehog target genes. We next used NIH3T3 Shh-Light II cells stably expressing a GLI-responsive luciferase reporter to further evaluate the effect of endosulfan on GLI transcriptional activity. Using the same treatment paradigm, we observed increased luciferase activity upon SAG treatment, whereas simultaneous treatment with SAG and endosulfan reduced luciferase activity twofold (**Figure EV3F**). Together, these results show that endosulfan downregulates hedgehog target gene expression.

Next, we tested whether endosulfan-mediated downregulation of *Gli* mRNAs was reflected in GLI protein levels. We first examined GLI3 levels in IMCD3 cells and found that SAG treatment stabilised GLI3 and increased the GLI3^FL^/GLI3^R^ ratio, whereas SAG–endosulfan cotreatment downregulated GLI3^FL^, significantly reducing the GLI3^FL^/GLI3^R^ ratio (**Figure 1F-G**). GLI3 accumulates at the ciliary tip upon hedgehog pathway activation [29]; in IMCD3 cells, we found that GLI3 accumulated at the ciliary tip upon SAG treatment, whereas SAG–endosulfan cotreatment significantly reduced this tip accumulation (**Figure 1H–I**). The reduced GLI3 ciliary tip accumulation is consistent with decreased GLI3^FL^ levels, since tip localisation is required for GLI3 activation. Consistent with impaired GLI3 activation, GLI1 levels, which are induced downstream of GLI3 upon hedgehog pathway activation, were stabilised by SAG treatment but reduced upon SAG–endosulfan cotreatment (**Figure 1J–K**). We further examined GLI1 levels in a parallel paradigm in which cells were treated with endosulfan after cilia formation. Even in this paradigm, GLI1 levels were stabilised upon SAG treatment and decreased upon cotreatment with SAG and endosulfan (**Figure EV2G–H**), indicating a robust effect of endosulfan on GLI downregulation.

Having established that endosulfan disrupts canonical hedgehog/GLI processing in ciliated kidney (IMCD3) and embryonic fibroblast (NIH3T3) cell lines, we asked whether this disruption also occurs in a tissue directly relevant to the clinical and epidemiological phenotypes of endosulfan exposure. Endosulfan exposure causes male infertility and reproductive defects [27], and hedgehog signalling is essential for spermatogenesis and Leydig cell function, prompting us to examine hedgehog target gene expression in the gonads of endosulfan-exposed mice. Transcriptomic analysis of bulk RNA-sequencing data from these gonads revealed downregulation of several hedgehog target genes, including *Gli1* and *Ptch1* (**Figure 2A and EV4A**), corroborating our *in cellulo* data. To validate these findings, we next used the mouse Leydig cell line TM3, which possesses primary cilia and responds to hedgehog signalling. At half the IC_50_ concentration (25.30 µM) (**Figure EV4B**), endosulfan did not affect ciliary number in TM3 cells (**Figure EV4C-D**), and the effects on ciliary length and IFT88 levels were consistent with those observed in IMCD3 and NIH3T3 cells (**Figure 2B-D**). Furthermore, SAG-mediated GLI3 ciliary tip accumulation and GLI1 stabilisation were both significantly reduced upon SAG–endosulfan cotreatment in TM3 cells (**Figure 2E-H**). Together, these results demonstrate that endosulfan suppresses hedgehog signalling by disrupting GLI3 processing and GLI1 expression across multiple cell types and in reproductive tissue.

**Figure 2.**
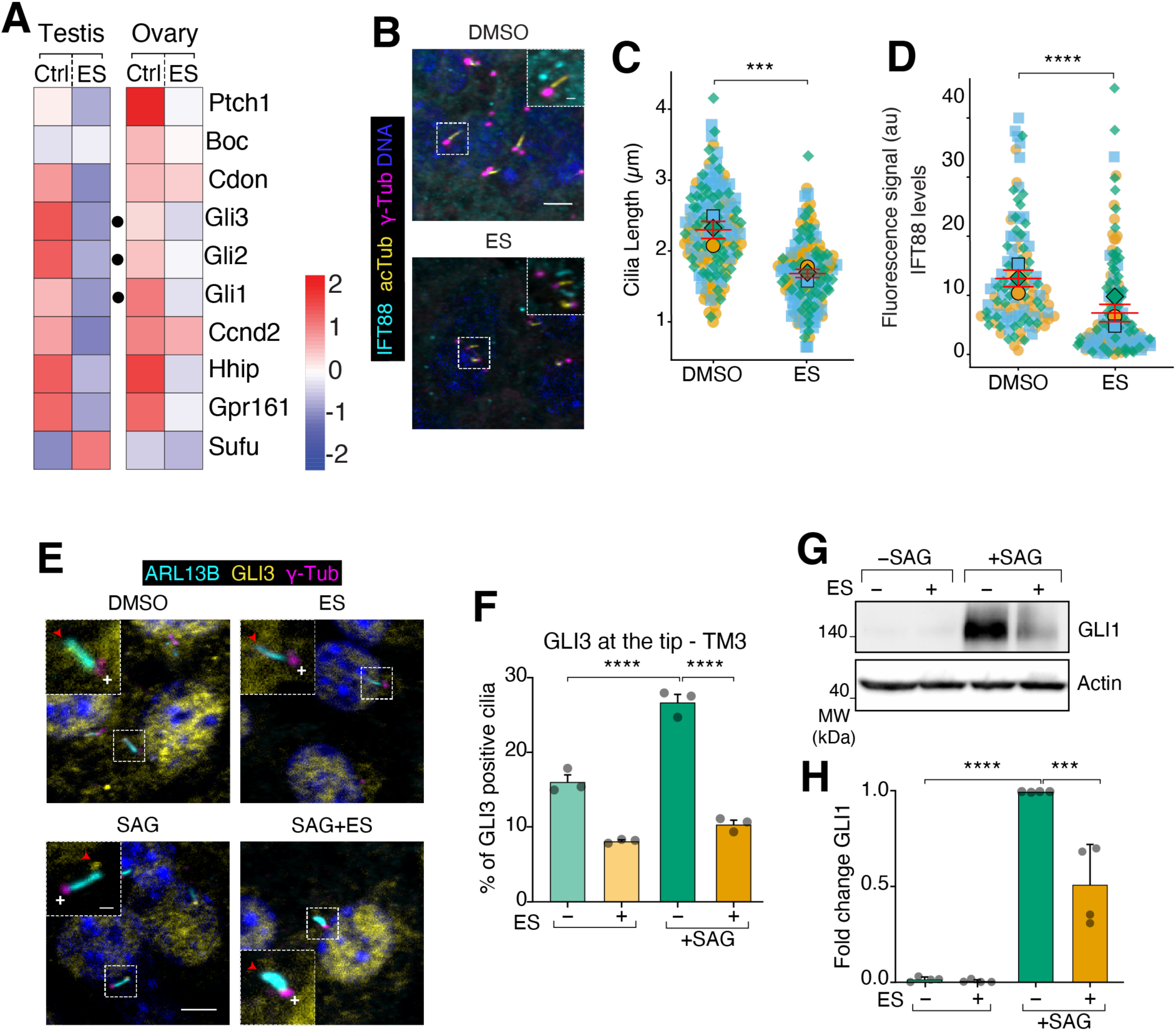
Endosulfan downregulates GLI expression and suppresses the Hedgehog pathway in gonads. **(A)** Heatmap of RNA-seq expression data for Hedgehog pathway genes in mouse gonads exposed to endosulfan. Gene expression values are shown as row-wise z-scores. **(B)** Mouse Leydig TM3 cells were treated with DMSO or endosulfan for 48 h, then fixed and stained for IFT88 (cyan), acTub (yellow), γ-Tubulin (magenta), and DNA (blue). Scale bars, 5 µm (main panel), 1 µm (inset). **(C-D)** Quantification of ciliary length **(C)** and IFT fluorescence intensities **(D)** for the cilia shown in **B**. Data represent all data points from three independent experiments, mean ±SEM. Cell numbers: acTub (DMSO n = 165, endosulfan n = 155) and IFT88 (DMSO n = 121, endosulfan n = 162). **(E)** TM3 cells were treated as in Figure 1E, fixed and stained for ARL13B (cyan), GLI3 (yellow), γ-Tubulin (magenta), and DNA (blue). Representative cilia are shown for each condition; the base of the cilia is marked with γ-Tubulin (shown with +), and the tip is marked with a red arrowhead. Scale bars, 5 µm (main panel), 1 µm (inset). **(F)**Cilia positive for GLI3 at the tip were quantified, and the percentage was plotted; data represent the mean ± SD of three independent experiments. **(G)** TM3 cells treated using the same paradigm as in Figure 1E were lysed, and the lysates were probed for GLI1 and actin (loading control) via immunoblotting. **(H)** Quantification of GLI1 levels. GLI1 signal intensities were measured and normalised against the loading control. GLI1 expression in the SAG-treated condition was set to 1. The experiment shown in **G** was repeated four times, and the data represent the mean ±SD of three independent experiments. All fluorescence intensities are expressed in arbitrary units (a.u.). Asterisks denote statistical significance as determined by unpaired Student’s t-test or one-way ANOVA followed by Tukey’s post hoc multiple comparison test, as appropriate: **** p ≤ 0.0001; *** p ≤ 0.001; ** p ≤ 0.01; * p ≤ 0.05.

### Endosulfan does not directly target hedgehog pathway components

We next sought to determine how endosulfan disrupts GLI processing and downregulates hedgehog signalling. The following are plausible mechanisms by which endosulfan can exert its effects: first, endosulfan competes with GLI, disrupting GLI-DNA interactions; second, endosulfan binds to PTCH1, SMO or GPR161, blocking their functions; third, endosulfan inhibits SAG-induced ciliary accumulation of SMO or GPR161 exit; fourth, endosulfan creates an imbalance in cAMP levels within the cilium; fifth, endosulfan binds to hedgehog pathway kinases – PKA and GSK3β – and rewires their activities (**Figure 3A**).

**Figure 3.**
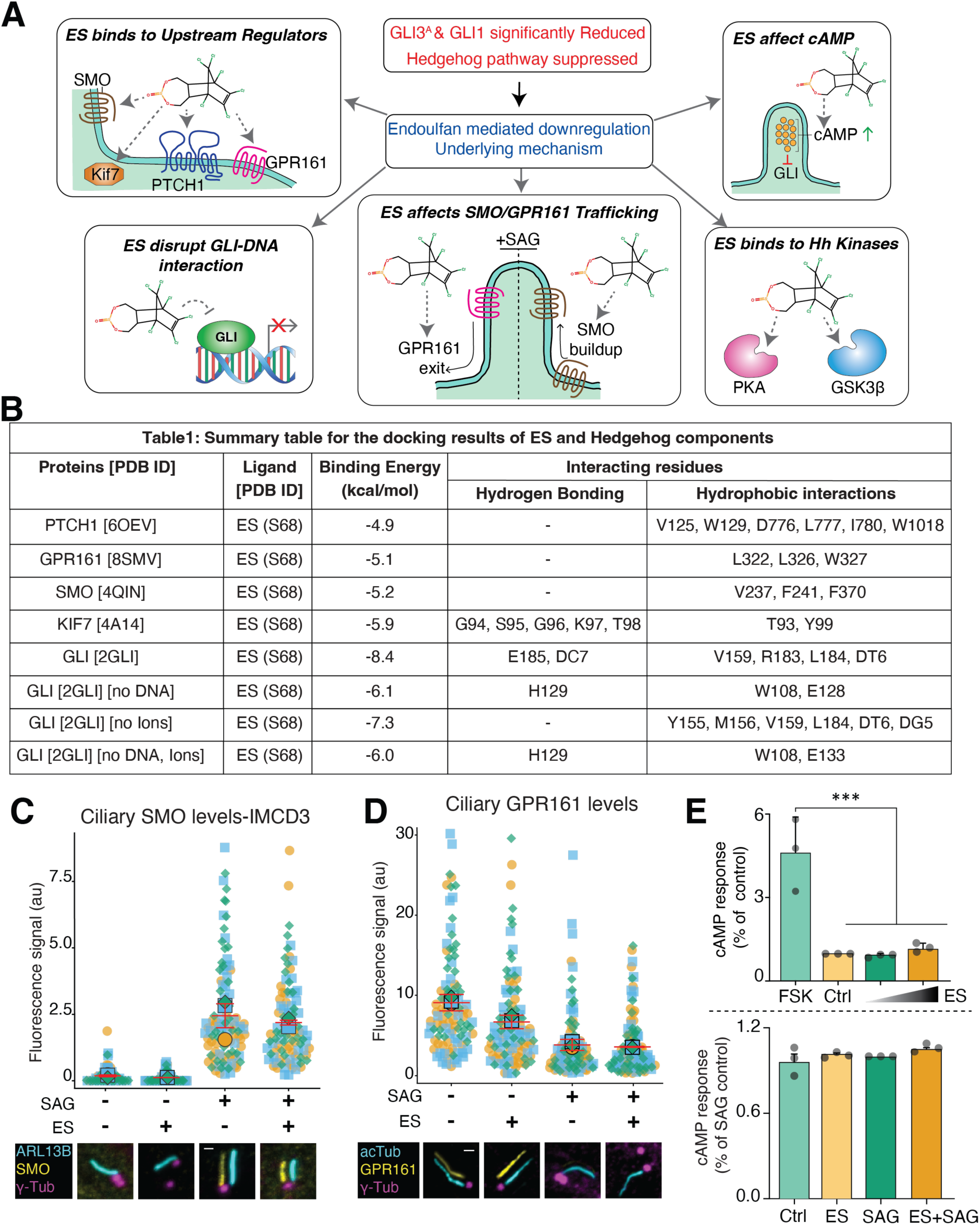
The effect of endosulfan on hedgehog signalling is independent of core pathway components. **(A)** Schematic illustration of hypothetical mechanistic models tested to identify the molecular target responsible for endosulfan-mediated downregulation of Hedgehog pathway gene expression. Model 1: endosulfan binds to the GLI–DNA complex, destabilising it and reducing target gene expression. Model 2: endosulfan binds upstream regulators of GLI, specifically PTCH1 or GPR161, to attenuate activation of the hedgehog pathway. Model 3: endosulfan inhibits SAG-induced SMO accumulation in the cilium and/or GPR161 trafficking out of the cilium. Model 4: endosulfan elevates intracellular cAMP levels, thereby attenuating formation of the activated GLI transcription factor. Model 5: endosulfan directly binds kinases that regulate GLI activity within the Hedgehog pathway. **(B)** Table 1 summarises the molecular docking results of endosulfan against hedgehog pathway components. **(C)** IMCD3 cells were treated as in Figure 1E, fixed and stained for ARL13B (cyan), SMO (yellow), and γ-Tubulin (magenta). Representative cilia are shown for each condition; the base of the cilia is marked with γ-Tubulin; corresponding full cell images are shown in **Figure EV6A**. Scale bar, 1 µm. The channels are shifted to facilitate visualisation of overlapping ciliary signals. Quantification of ciliary SMO fluorescence intensities. In the superplots, three independent experimental data points are represented by three different shapes, with colours indicated. Each square, circle, or triangle represents a data point, and the larger version of its shape represents the mean. Data represent all data points from three independent experiments, mean ±SEM. Cell counts per condition: DMSO, n = 104; endosulfan, n = 129; SAG, n = 106; SAG + endosulfan, n = 95. **(D)** IMCD3 cells stably expressing GPR161 fused to the fluorescent protein NG at its intracellular C-terminus were treated as in Figure 1E. Representative immunofluorescence micrographs show acetylated tubulin (cyan), GPR161 (yellow) imaged via its intrinsic NG fluorescence, and γ-tubulin (magenta); corresponding full cell images are shown in **Figure EV6F**. Scale bar, 1 µm. Superplot convention as in C. Cell counts per condition: DMSO, n = 100; endosulfan, n = 106; SAG, n = 103; endosulfan + SAG, n = 115. **(E)** Quantification of intracellular cAMP levels in IMCD3 cells. In one setup, cells were treated with forskolin, DMSO, or endosulfan (12.5 µM). The cAMP level in the vehicle-treated control (DMSO) condition was set to 1 and compared with the other conditions. In another setup, forskolin was added to all conditions to elevate basal cAMP levels, and cells were then treated with endosulfan alone or in combination with SAG. The cAMP level in the SAG condition was set to 1. Data represent the mean ±SD of three independent experiments. Fluorescence intensity is expressed in arbitrary units (a.u.). Asterisks denote statistical significance as determined by one-way ANOVA followed by Tukey’s post-hoc multiple comparison test: **** p ≤ 0.0001; *** p ≤ 0.001; ** p ≤ 0.01; * p ≤ 0.05.

Next, we tested each of these possibilities. We employed *in silico* molecular docking followed by molecular dynamics (MD) simulations to assess endosulfan binding to PTCH1, SMO, GPR161, KIF7, and GLI (**Figure 3B** and **Figure EV5A-G**). First, we performed molecular docking of endosulfan at the active sites of PTCH1, SMO, and GPR161. The binding energies for each were below the -5.5 kcal/mol cut-off we set. We did not proceed with MD simulations and discarded the possibility of endosulfan binding directly to them. The KIF7-endosulfan binding energy was -5.9 kcal/mol at its ATP-binding site; therefore, we proceeded with MD simulations, which showed that endosulfan did not bind KIF7 stably (**Movie EV1**). Next, we assessed whether endosulfan binds directly to GLI or to the GLI-DNA complex. GANT61, a GLI1 antagonist, docks between ZF-2 and ZF-3, either in the presence or absence of DNA [30,31]. For endosulfan, we asked whether it binds between ZF-4 and ZF-5 to inhibit GLI1-DNA interactions, or docks between ZF-2 and ZF-3, as with GANT61. To address this, we performed molecular docking and MD simulations. We observed that endosulfan does not bind to the DNA-facing ZF-4 and ZF-5 pockets, but docks between ZF-2 and ZF-3 (**Figure EV5H**). Thereafter, we ran an MD simulation to probe binding. In a 400 ns simulation, endosulfan remained stable within the binding pocket for approximately 130 ns, after which progressive destabilisation, reflected in increasing protein RMSD (data not shown), led to complete ligand exit by ∼290 ns (**Movie EV2**), confirming the unstable binding of endosulfan with the GLI-DNA complex. We next examined the effect of endosulfan on ciliary trafficking of SMO and GPR161. SMO accumulates in primary cilia upon SAG treatment; however, endosulfan did not affect SAG-induced SMO accumulation in primary cilia in IMCD3, NIH3T3, and TM3 cells (**Figure 3C** and **EV6B-E**). GPR161 exits cilia upon hedgehog pathway activation; we found that endosulfan did not affect SAG-dependent ciliary exit (**Figure 3D**). SMO and GPR161 coordinate ciliary cAMP levels, which are crucial for GLI^FL^ formation upon pathway activation. We next tested whether endosulfan affects cAMP levels; the cAMP measurements showed that endosulfan neither increased cAMP levels nor reduced forskolin-mediated increases in cAMP (**Figure 3E**). Together, these results suggest that endosulfan exposure does not modulate the localisation and/or function of hedgehog pathway receptors or GLI.

### Endosulfan engages PKA and GSK3β through distinct allosteric mechanisms

PKA phosphorylates GLI2 and GLI3 at specific serine residues, triggering sequential phosphorylation by GSK3β, promoting GLI2/GLI3 interaction with βTrCP, and generating the transcriptional repressor forms GLI2^R^ and GLI3^R^ [6,7]. We assessed whether endosulfan rewires PKA and GSK3β to exert its effect on GLI processing. We performed molecular docking and MD simulations of endosulfan with the PKA catalytic subunit (PKA-C) and GSK3β.

The MD simulation of the PKA-C-endosulfan complex over 400 ns revealed key interactions that underpin endosulfan’s stability and affinity within the protein’s binding pocket (**Figure 4A**). These interactions include hydrophobic interactions, hydrogen bonding, and halogen bonding (**Figure EV7A**). Each plays a distinct and crucial role in maintaining endosulfan’s binding orientation and overall stability. The surface potential analysis of PKA-C upon endosulfan binding revealed no change in surface potential at the binding site in comparison to the apo form (**Figure 4B** and **EV7B**). The stability of PKA-C in the presence of endosulfan was evaluated using the RMSD of the backbone atoms and the compactness of the protein backbone, as measured by Rg values, during MD simulations. The conservation of RMSD and Rg indicates the absence of major structural perturbation (**Figure 4C**). The RMSF of backbone residues was compared between PKA-C-apo and PKA-C-endosulfan. We observed a significant reduction in RMSF in two regions upon endosulfan binding: the G-rich loop and the activation loop. The near-superimposable RMSF profile demonstrates that the endosulfan effect on protein dynamics is restricted to the G-rich loop and the activation loop (**Figure 4D** and **EV7C**). The relative solvent-accessible surface area (rSASA) of PKA-C-endosulfan shows no change in comparison to PKA-C-apo. In ΔrSASA, the signal oscillates symmetrically around zero across the entire protein (residues 19–351), with the vast majority of values within ±0.05%. (**Figure 4E-F** and **EV7D**). The free energy landscape (FEL), constructed using RMSD and Rg as collective variables, showed an identical global minimum for both the apo and endosulfan-bound states (**Figure EV7E**). To assess whether endosulfan induces any conformational change in PKA-C, the differential free energy landscape (ΔFEL = G_drug-bound_ − G_apo_) was computed using RMSD and Rg as collective variables. The negligible ΔFEL between the endosulfan-bound and apo states indicates that endosulfan does not induce major global conformational change (**Figure 4G**) and conformational entropy (**Figure EV7F**), consistent with its effect being restricted to localised dynamics at the G-rich loop and activation loop. To further understand the effect of endosulfan binding on PKA activity, we performed an *in vitro* kinase assay, which showed a modest increase in PKA activity in the presence of endosulfan (**Figure 4H**). Together, these results suggest that endosulfan increases PKA activity without grossly perturbing the protein; however, it subtly restricts motions in the G-rich loop and activation loop, a classic mechanism by which ligands allosterically tune kinase activity.

**Figure 4.**
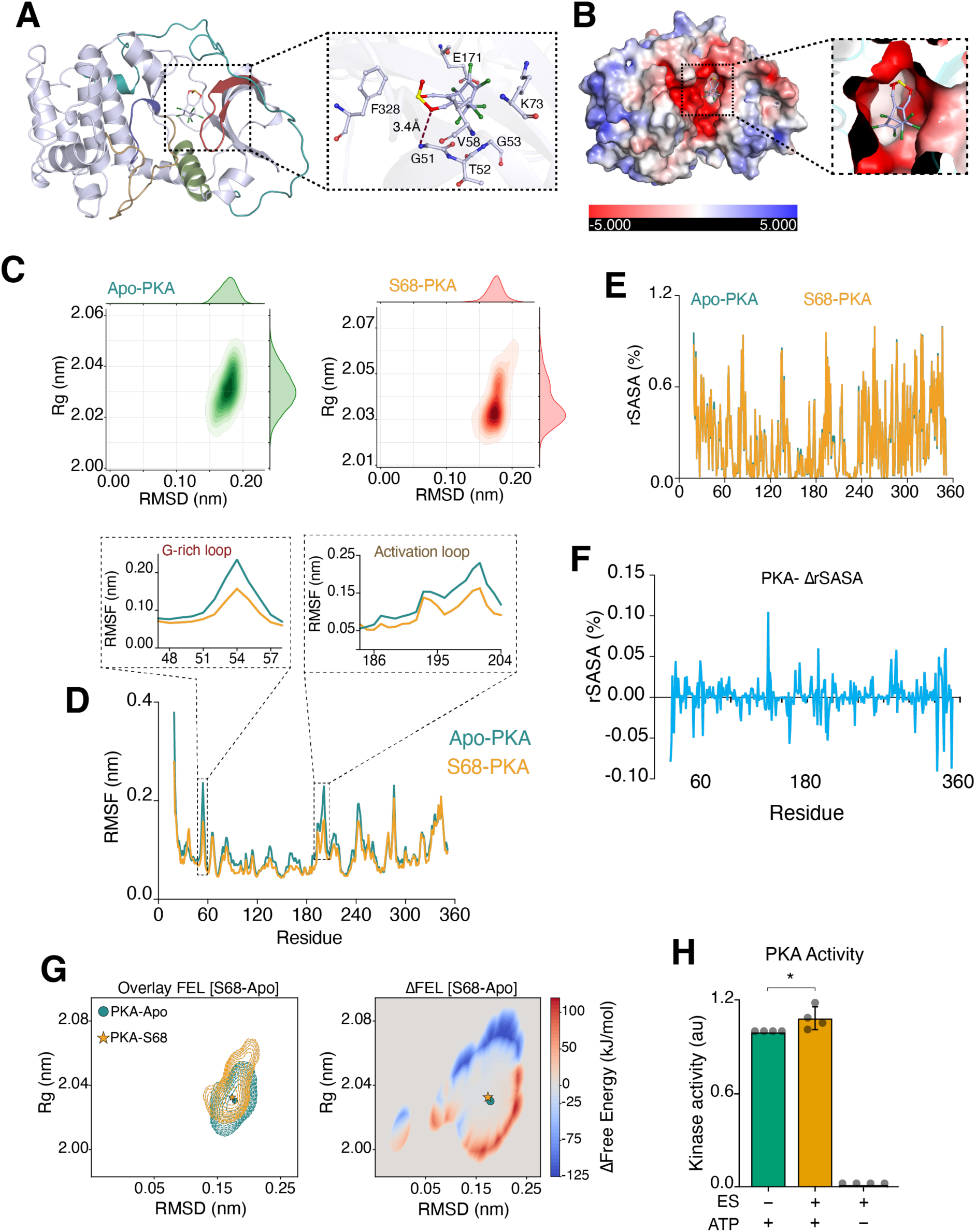
Endosulfan binds and hyperactivates the PKA catalytic subunit. **(A)** Structure of the PKA-C: endosulfan complex after a 400 ns MD simulation, highlighting the ATP-binding site (green) and substrate-binding site (cyan). The inset shows key residues involved in interactions with endosulfan throughout the 400 ns simulation; hydrogen bonds are depicted as dashed lines (Å). **(B)** Electrostatic surface potential representation of the PKA-C: endosulfan complex. The inset shows a zoomed view of the electrostatic surface potential at the endosulfan binding site. **(C)** RMSD versus radius of gyration (Rg) plot of PKA-C in the apo (green) and endosulfan-bound (red) forms, demonstrating the structural stability and compactness of PKA-C throughout the 400 ns MD simulation. **(D)** Root mean square fluctuation (RMSF) of PKA-C residues in the apo (green) and endosulfan-bound (wheat) forms. The inset shows the RMSF of the G-rich loop (residues 47–58) and the activation loop (residues 184–204). **(E)** Relative solvent-accessible surface area (rSASA) expressed as a percentage for individual PKA-C residues, illustrating changes in solvent accessibility between the apo and endosulfan-bound forms. **(F)** Delta relative solvent-accessible surface area (ΔrSASA) expressed as a percentage for individual PKA-C residues, illustrating changes in solvent accessibility. **(G)** Two-dimensional overlay and change in free energy landscape (ΔFEL) of PKA-C in the apo and endosulfan-bound forms. The global minimum energy state in the apo form is indicated by a filled circle (●) and in the endosulfan-bound form by a filled star (★). **(H)** Quantification of *in vitro* PKA kinase activity in the presence of endosulfan. Kinase activity in the absence of endosulfan was set to 1. Data represent the mean ±SD from four independent experiments. Kinase activity is expressed in arbitrary units (a.u.). The MD simulations and the corresponding analyses were performed in triplicate, and the representative graphs show the mean across three independent simulations for each parameter analysed. Asterisks denote statistical significance as determined by one-way ANOVA followed by Tukey’s post-hoc multiple comparison test: **** p ≤ 0.0001; *** p ≤ 0.001; ** p ≤ 0.01; * p ≤ 0.05.

We next tested whether endosulfan binds to and modulates GSK3β activity. The 500 ns MD simulation of the GSK3β-endosulfan complex revealed key interactions that underpin endosulfan’s stability and affinity within the GSK3β binding pocket (**Figure 5A**). These interactions include hydrophobic interactions, hydrogen bonding, and halogen bonding (**Figure EV8A**). Similar to the surface potential analysis of PKA-C, in GSK3β, upon endosulfan binding, no change in surface potential was observed at the binding site in comparison to the apo form (**Figure 5B** and **EV8B**). The RMSD vs Rg plots for both conditions share an identical global minimum Rg (∼2.121 nm) but differ in the global minimum RMSD: ∼0.205 nm for the apo state versus ∼0.175 nm for the endosulfan-bound state (**Figure 5C**). The shift in RMSD with a constant Rg is a hallmark of conformational state selection without gross structural change. We compared the RMSF of backbone residues between GSK3β-apo and GSK3β-endosulfan. The trough is broad, spanning approximately 40 residues, not just at the peak residue, confirming that the ordering effect encompasses the entire activation loop mid-segment, not just one or two residues. We observed a significant reduction in RMSF in the activation loop at residue 210: ∼0.29 nm GSK3β-apo vs ∼0.20 nm GSK3β-endosulfan (**Figure 5D** and **EV8C**). The near-superimposable RMSF profile indicates that endosulfan’s effect on protein dynamics is restricted to the activation loop. Despite significant regional differences in activation-loop dynamics, the total SASA (**Figure EV8D**) and rSASA of GSK3β-endosulfan show no change compared with GSK3β-apo (**Figure 5E**). In ΔrSASA, residue ∼62 shows the largest single-residue deviation (+0.155%, with a sharp negative companion at ∼residue 65, reaching −0.10%). This sharp positive-then-negative pattern at adjacent residues is characteristic of a local conformational reorientation rather than genuine burial or exposure. The activation loop region (∼residues 200–240) shows the most sustained and structured deviation. There is a notable negative excursion reaching −0.10% at ∼residue 213, flanked by a positive peak at ∼180 and a deeper negative trough at ∼residue 235 (−0.13%). The negative ΔrSASA at these positions means the drug-bound protein has less solvent exposure at specific activation loop residues (**Figure 5F**). The ΔFEL shows that the endosulfan-bound state preferentially stabilises the active conformation (RMSD ∼0.10–0.17 nm, Rg ∼2.121 nm), reaching energies of −75 to −100 kJ/mol, whereas the apo state preferentially stabilises the inactive conformation (RMSD ∼0.17–0.27 nm, Rg ∼2.121 nm), reaching +100 to +125 kJ/mol (**Figure 5G** and **EV8E**). Thus, the ΔFEL indicates that endosulfan binding induces conformational changes in GSK3β despite changing the conformational entropy (**Figure EV8F**). Together, these analyses suggest that, unlike PKA-C, where endosulfan binding left the conformational landscape largely unaltered, GSK3β shows a clear endosulfan-induced shift towards the active-state population, consistent with a conformational selection mechanism.

**Figure 5.**
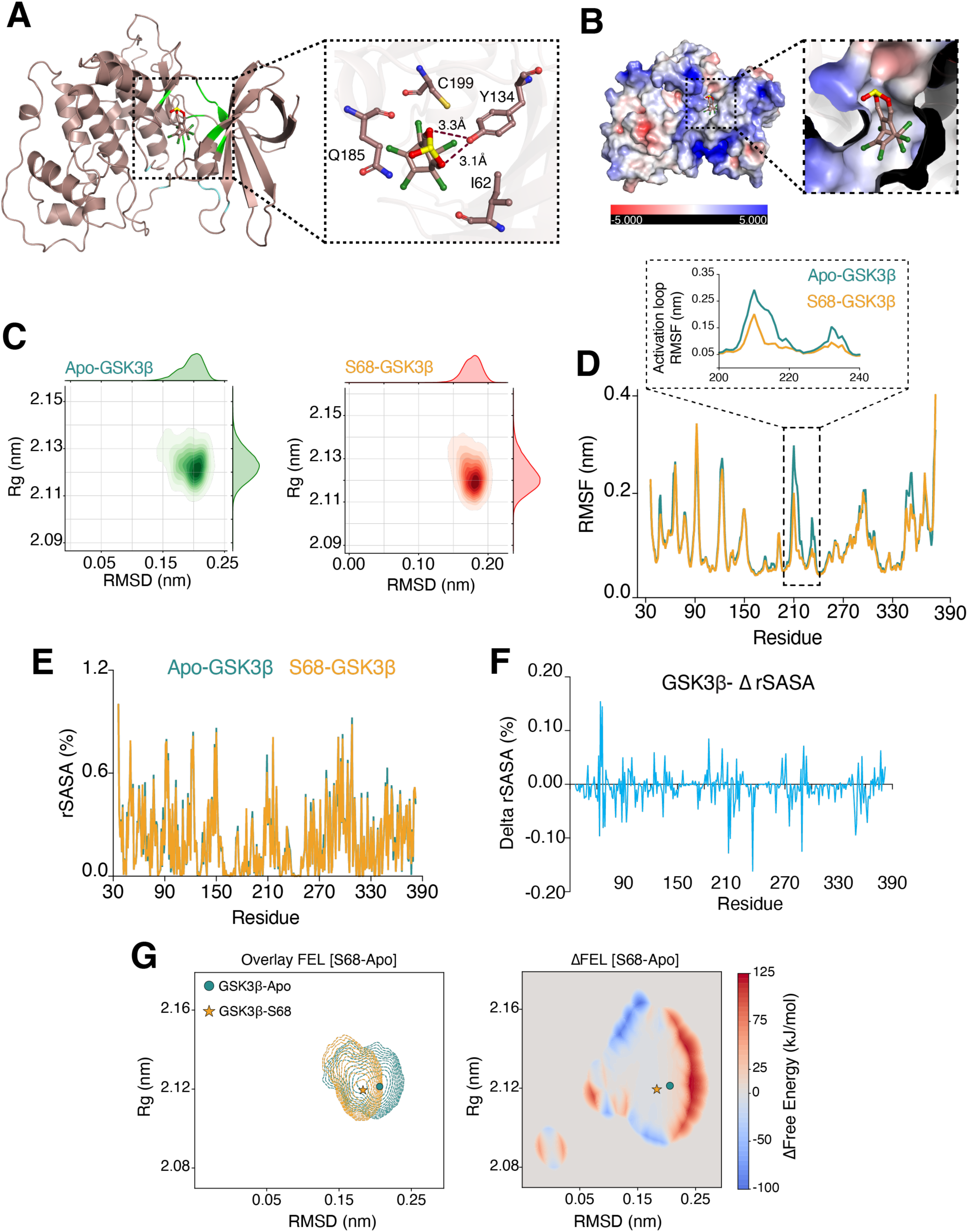
Endosulfan directly induces conformational changes in GSK3β and stabilises its active state. **(A)** Representative structure of the GSK3β: endosulfan complex after 500 ns MD simulation, highlighting the ATP-binding site (green) and substrate-binding site (cyan). The inset shows key residues involved in interactions with endosulfan throughout the 500 ns simulation; hydrogen bonds are depicted as dashed lines in (Å). **(B)** Electrostatic surface potential representation of the GSK3β: endosulfan complex. The inset shows a zoomed view of the electrostatic surface potential at the endosulfan binding site. **(C)** RMSD versus radius of gyration (Rg) plot of GSK3β in the apo (green) and endosulfan-bound (red) forms, demonstrating the structural stability and compactness of GSK3β throughout the 500 ns MD simulation. **(D)** Root mean square fluctuation (RMSF) of GSK3β residues in the apo (green) and endosulfan-bound (wheat) forms. The inset shows the RMSF of the activation loop (residues 200–240). **(E)** Relative solvent-accessible surface area (rSASA) expressed as a percentage for individual GSK3β residues, illustrating changes in solvent accessibility between the apo and endosulfan-bound forms. **(F)** Delta relative solvent-accessible surface area (ΔrSASA) expressed as a percentage for individual PKA-C residues, illustrating changes in solvent accessibility. **(G)** Two-dimensional overlay and change in free energy landscape (ΔFEL) of GSK3β in the apo and endosulfan-bound forms. The global minimum energy state in the apo form is indicated by a filled circle (●) and in the endosulfan-bound form by a filled star (★). The MD simulations and the corresponding analyses were performed in triplicate, and the representative graphs show the mean across three independent simulations for each parameter analysed.

### Endosulfan hyperactivates GSK3β to disrupt GLI processing

To further validate the *in silico* GSK3β-endosulfan binding, we performed the Cellular Thermal Shift Assay (CETSA) on GSK3β in the presence of endosulfan. CETSA analysis confirms the direct binding between GSK3β and endosulfan. However, it also suggests that endosulfan binding decreases the aggregation temperature (T_agg_) of GSK3β (**Figure 6A–B**). This is consistent with our MD simulations, in which endosulfan binding shifted the conformational equilibrium toward an active state that may be less thermally stable than the apo-enriched inactive state, providing a structural explanation for the observed decrease in T_agg_. A negative thermal shift is a recognised signature of target engagement. As established in the context of CETSA, the thermal response is governed not only by ligand affinity but also by the thermodynamics and kinetics of the entire protein-ligand system, including the relative stability of the conformational states to which the ligand preferentially binds [32]. The recently established DeepKinomeWeb server provides quantitative predictions of binding affinity to evaluate kinase–inhibitor interactions. We compared the predicted binding affinities of endosulfan and the known GSK3β inhibitors BRD3731 and staurosporine against GSK3β to determine whether endosulfan is more likely to act as an inhibitor or an activator. Further, superposition of the GSK3β–staurosporine and GSK3β–endosulfan complexes revealed that the two ligands occupy distinct binding sites (**Figure 6C**), supporting the conclusion that endosulfan does not act as a classical ATP-competitive inhibitor of GSK3β. Together, the negative CETSA thermal shift and the increased kinase activity provide direct evidence that endosulfan engages GSK3β in a manner that shifts its conformational equilibrium towards a less thermally stable, more active state, consistent with the conformational selection mechanism identified by MD simulation.

**Figure 6.**
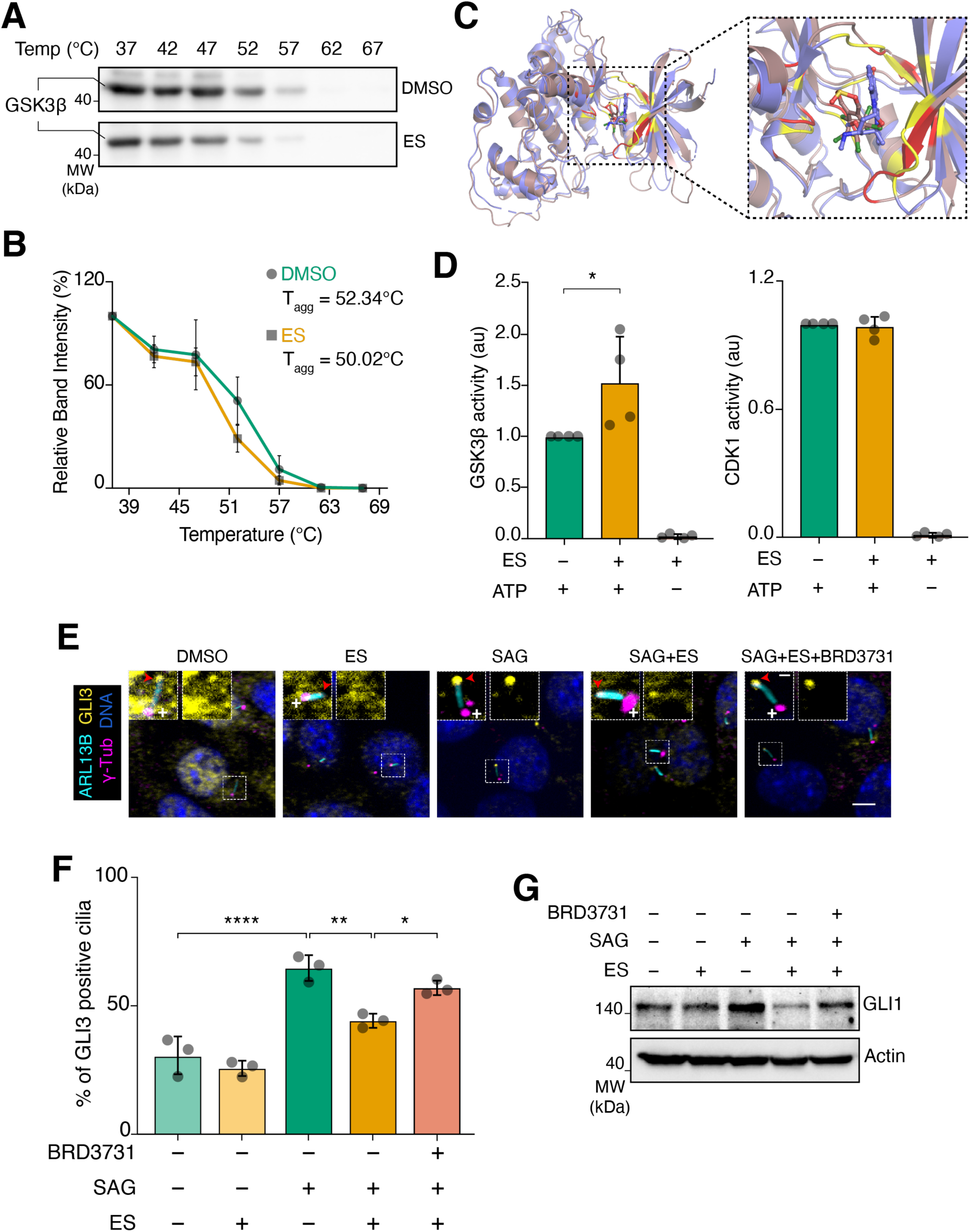
Characterisation of GSK3β as a direct endosulfan target and its role in hedgehog pathway regulation. **(A)** IMCD3 cells were treated with DMSO or endosulfan for 1 h, followed by a 3-minute incubation at temperatures ranging from 37 ℃ to 67 ℃ in increments of 5 ℃. Cells were lysed, and the lysates were immunoblotted for GSK3β. **(B)** Quantification of GSK3β protein stability. Signal intensities for GSK3β were measured and plotted. GSK3β expression at 37 ℃ in both conditions was set to 100%. Data represent the mean ±SD of four independent experiments. **(C)** Superimposed structures of staurosporine-bound (purple; PDB: 1Q3D) and endosulfan-bound (brown) GSK3β, illustrating the binding regions of staurosporine (yellow) and endosulfan (red) at the GSK3β active site, showing that these molecules do not occupy the same binding region. **(D)** Quantification of *in vitro* GSK3β and CDK1/CyclinA2 kinase activity in the presence of endosulfan. Kinase activity in the absence of endosulfan was set to 1. Data represent the mean ± SD of four independent experiments. Kinase activity is expressed in arbitrary units (a.u.). **(E)** IMCD3 cells were treated with the given conditions, fixed and stained with ARL13B (cyan), GLI3 (yellow), γ-tubulin (magenta), and DNA (Blue). The base of the cilia is marked with γ-tubulin denoted by + and the tip is marked with a red arrowhead. Scale bars, 5 µm; inset scale bars, 1 µm. **(F)** Quantification of the percentage of cilia positive for GLI3 tip accumulation **E**. Quantifications were performed similarly to Figure 1I. Data represent the mean ±SD of three independent experiments. **(G)** Quantification of GLI1 levels. IMCD3 cells treated with BRD3731, in the presence or absence of endosulfan and/or SAG, were lysed and immunoblotted for GLI1. Signal intensities for GLI1 were measured and plotted. Data represent the mean ±SD of four independent experiments. Asterisks denote statistical significance as determined by one-way ANOVA followed by Tukey’s post-hoc multiple comparison test: **** p ≤ 0.0001; *** p ≤ 0.001; ** p ≤ 0.01; * p ≤ 0.05.

Next, we sought to test whether the effects of endosulfan on the hedgehog pathway could be rescued by the known GSK3β inhibitor BRD3731. We assessed GLI3 accumulation at the tips of primary cilia upon SAG treatment in IMCD3 cells. SAG treatment led to accumulation of GLI3 at the tips of cilia, which was significantly reduced upon co-treatment with endosulfan. However, we found that the addition of BRD3731 partially rescued the GLI3 tip accumulation caused by endosulfan treatment (**Figure 6E-F**). Furthermore, we assessed total GLI1 levels and found that SAG stabilised GLI1, which decreased upon co-treatment with SAG and endosulfan. However, the addition of BRD3731 rescued this decrease, bringing GLI1 levels back to control levels (**Figure 6G**). Together, these results suggest that endosulfan binds to and hyperactivates GSK3β, disrupting GLI processing, a phenotype that GSK3β inhibitors can rescue.

### Endosulfan selectively targets GSK3β over structurally related kinases

Having established that endosulfan activates both PKA and GSK3β, we next asked whether this represented a selective effect on specific Hedgehog pathway kinases or a more general perturbation of protein kinase activity. To address this, we first examined the predicted binding profile of endosulfan across a panel of 225 kinases using DeepKinomeWeb, which generates quantitative binding affinity predictions for this kinase panel [57]. Strikingly, endosulfan’s predicted profile across the 225-kinase panel was qualitatively distinct from all five reference kinase inhibitors examined - BRD3731, staurosporine, ponatinib, dasatinib, and sorafenib which showed broad inhibitory signatures across the panel. By contrast, endosulfan received predominantly activator-range predictions across most kinases, with GSK3β among the highest-scoring kinases in this category **(Figure EV9)**. This unique activator-range profile across the kinome suggested that endosulfan’s effect on GSK3β activity is not a generic consequence of kinase active-site engagement.

To directly test kinase selectivity experimentally, we examined the effect of endosulfan on CDK1. This structurally related kinase shares the conserved bilobal kinase fold and DFG motif with GSK3β and plays a central role in cell cycle progression. We performed an *in vitro* kinase assay using purified CDK1–Cyclin A2 complex in the presence of endosulfan. In contrast to the significant increase in GSK3β activity observed under the same conditions, endosulfan did not affect CDK1 kinase activity **(Figure 6D)**. To further examine the structural basis of this selectivity, we performed molecular dynamics simulations of endosulfan docked to CDK1 (PDB ID: 6GU2) and CDK2 (PDB ID: 1H1P) for 500 ns each. Unlike the stable, persistent interactions observed in the GSK3β–endosulfan complex over 500 ns, endosulfan did not form a stable complex with either CDK1 or CDK2 over the course of the simulation, with progressive ligand destabilisation and exit from the binding pocket observed in both cases **(Movie EV5 and EV6)**. Together, these results demonstrate that endosulfan’s kinase-activating effect is selective for GSK3β and does not represent a generalised perturbation of kinase activity, consistent with the distinct structural features of the GSK3β binding site identified by MD simulation and the non-overlapping binding site relative to the ATP-competitive inhibitor staurosporine **(Figure 6C)**.

### Endosulfan represses GLI1-regulated ciliary and centrosomal genes

A canonical view of hedgehog pathway regulation is as follows: IFT machinery maintains cilia, which enable hedgehog signalling via GLI processing and activation, and in turn control hedgehog-dependent expression of GLI target genes [33]. Several ciliary-centrosomal genes govern hedgehog signalling in a feedback-loop manner [34,35]. Hence, we next sought to identify novel GLI target genes associated with centrosomes-cilia and hedgehog signalling.

Using the MEME Suite, we analysed GLI consensus binding motifs across 241 cilia-related genes and identified 41 candidate GLI target genes. In parallel, using GLI1 ChIP-seq data, we independently identified GLI1 binding sites in the promoters of 55 genes, drawn from a set of 99 genes comprising 31 Hedgehog pathway genes and 68 differentially expressed ciliary genes identified by RNA-seq. FIMO analysis of these 55 genes identified 14 genes harbouring a GLI1 binding motif. Comparing the MEME-derived (41 genes) and ChIP-seq/FIMO-derived (14 genes) candidate sets identified *Cetn3* (CETN3/Centrin3) as the only gene shared by both analyses (**Figure 7A** and **EV10A-B**). We validated *Cetn3* as a hedgehog-responsive gene by qRT-PCR: *Cetn3* expression was significantly upregulated upon SAG treatment, whereas co-treatment with SAG and endosulfan led to a significant decrease in Cetn3 expression (**Figure 7B**). Consistently, CETN3 protein levels at the ciliary base were upregulated upon SAG treatment and significantly reduced upon SAG–endosulfan cotreatment (**Figure 7C-D** and **EV11A**).

**Figure 7:**
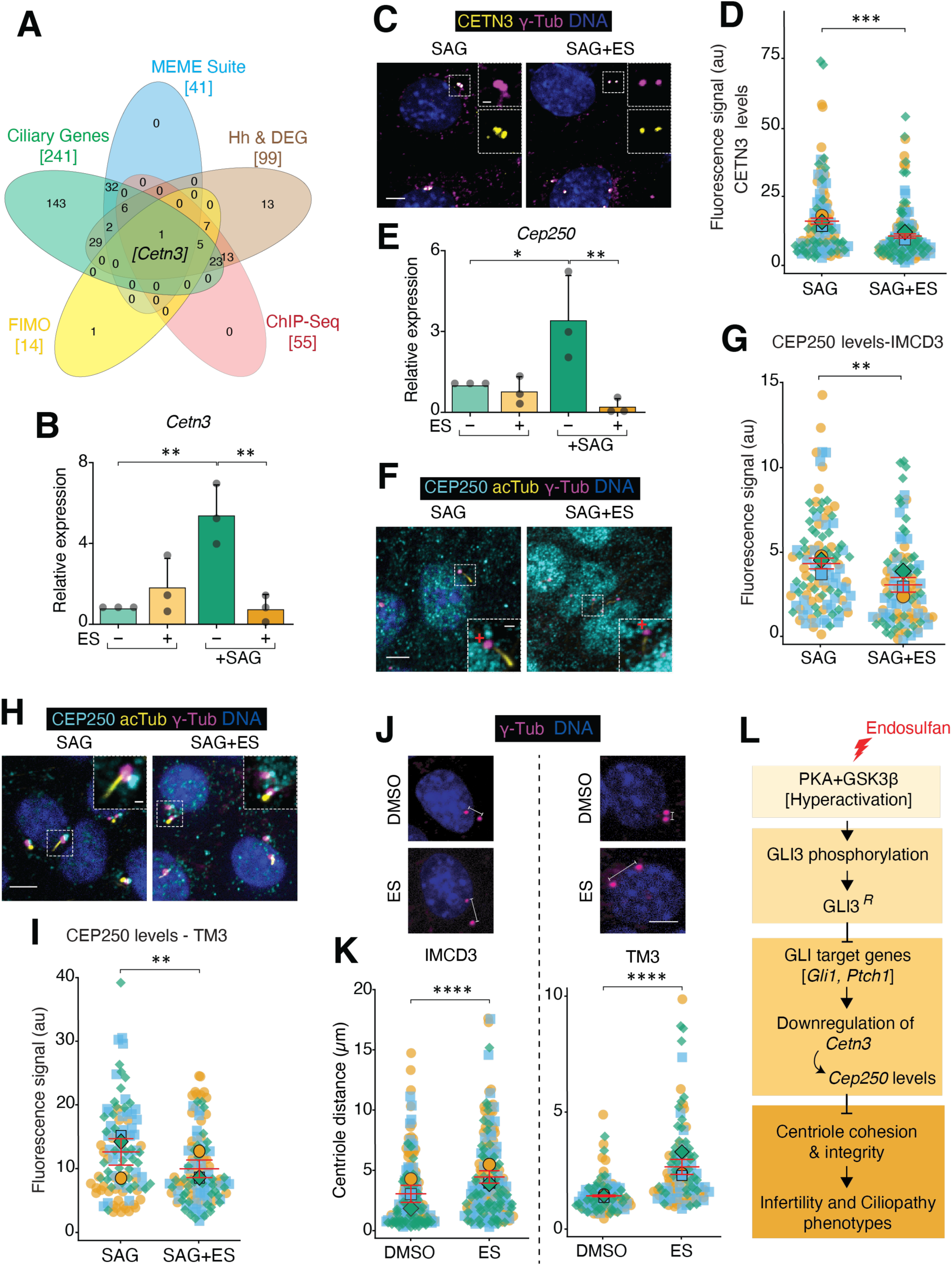
Endosulfan perturbs the expression of centrosomal genes downstream of GLI1. **(A)** A Venn diagram showing the overlap between candidate GLI target genes identified by MEME Suite motif analysis and ChIP-seq analysis across 241 ciliary genes. The analysis reveals *Cetn3* as a potential GLI target. **(B)** qRT-PCR analysis to validate *Cetn3* as a GLI target gene. IMCD3 cells were treated as indicated, and mRNA expression was analysed. Gene expression was normalised to the DMSO condition (set to 1). Data represent the mean ± SD of three independent experiments. **(C)** and **(D)** Representative immunofluorescence micrographs showing CETN3 (yellow), γ-tubulin (magenta), and DNA (Blue) **(C)** and quantification of CETN3 levels at the ciliary base **(D)** in IMCD3 cells treated with SAG in the presence or absence of endosulfan. Data represent all data points from three independent experiments mean ± SEM. Cell numbers: SAG n = 112, SAG + endosulfan n = 151. **(E)** qRT-PCR analysis of *Cep250* in IMCD3 cells treated as indicated. Gene expression was normalised to the DMSO condition (set to 1). Data represent the mean ± SD of three independent experiments. **(F)** IMCD3 cells were treated as indicated, and representative immunofluorescence micrographs show CEP250 (cyan), acTub (yellow), γ-tubulin (magenta), and DNA (Blue). **(G)** Quantification of CEP250 levels at the ciliary base. Superplot convention as in Figure 1B. Cell numbers: SAG n = 107, SAG + endosulfan n = 105. **(H)** TM3 cells were treated as indicated, and representative immunofluorescence micrographs show CEP250 (cyan), acTub (yellow), γ-tubulin (magenta), and DNA (Blue). **(I)** Quantification of CEP250 levels at the ciliary base. Superplot convention as in Figure 1B. Cell numbers: SAG n = 102, SAG + endosulfan n = 110. **(J)** Representative immunofluorescence micrographs of IMCD3 and TM3 cells treated as indicated, showing γ-tubulin (magenta) and DNA (Blue). **(K)** Quantification of inter-centriolar distance measured from IMCD3 and TM3 cells. Superplot convention as in Figure 1B. Cell numbers: for IMCD3-FlpIn cells, DMSO n = 179, endosulfan n = 180; for TM3 cells, DMSO n = 108, endosulfan n = 109. **(L)** Schematic diagram showing the mechanism of action of endosulfan via PKA and GSK3β to regulate centriole cohesion and integrity, leading to infertility and ciliopathies. Fluorescence intensity is expressed in arbitrary units (a.u.). Scale bars, 5 µm; inset scale bars, 1 µm. Asterisks denote statistical significance as determined by unpaired Student’s t-test or one-way ANOVA followed by Tukey’s post-hoc multiple comparison test, as appropriate: **** p ≤ 0.0001; *** p ≤ 0.001; ** p ≤ 0.01; * p ≤ 0.05.

A recent study showed that Centrin3 plays a crucial role in the microtubule-organising centre and regulates RNA splicing of centriole-related genes, including *Cep250* [36]. Building on this link, we asked whether *Cep250* expression is regulated by the GLI-*Centrin3* axis and whether endosulfan would disrupt it. We found that *Cep250* expression was controlled in a hedgehog-dependent manner: SAG treatment increased *Cep250* levels, whereas co-treatment with SAG and endosulfan significantly reduced its expression (**Figure 7E**). Further indirect immunofluorescence analysis revealed that in both IMCD3 and TM3 cells, CEP250 levels at the base of the cilium were significantly reduced in the presence of endosulfan (**Figure 7F-I and EV11B-C**). Earlier reports indicate that CEP250 mutations or deletions affect centriole cohesion [37,38]. So, we measured the distance between centrioles in the presence of endosulfan and observed a significant increase in this distance in both IMCD3 and TM3 cells (**Figure 7J-K**). Together, these results systematically demonstrate that *Cetn3* is a GLI target, and that *Cep250* downstream of *Cetn3* can be regulated by GLI and hedgehog signalling (**Figure 7L**). Moreover, these results provide the first evidence that endosulfan could disrupt this positive feedback axis.

## Discussion

Here we show that the organochlorine insecticide endosulfan disrupts primary cilium-dependent hedgehog signalling through a mechanism distinct from all previously described pathway antagonists: rather than acting at the level of ciliary receptors, cAMP production, or GLI–DNA binding, endosulfan directly rewires the activity of two core pathway kinases, PKA and GSK3β, suppressing GLI-dependent transcription and the expression of newly identified GLI target genes required for ciliary and centrosomal integrity. Pesticide exposure is associated with a substantial human health burden, including congenital disabilities, skeletal anomalies, and reproductive failure [23,25], and endosulfan itself induces genotoxicity via reactive oxygen species in a time- and concentration-dependent manner [26]. Disruption of hedgehog signalling produces a remarkably overlapping set of phenotypes, including dorsalisation defects, impaired cerebral growth, loss of left–right asymmetry, skeletal anomalies, and infertility [39,40], raising the possibility that suppression of the hedgehog pathway contributes to the developmental consequences of endosulfan exposure alongside its established neurotoxicity. Establishing this causal link definitively will require genetic rescue experiments and appropriate animal models in which endosulfan-induced developmental phenotypes are rescued by restoring hedgehog pathway activity an important direction for future work.

Before examining pathway-specific effects, we confirmed that endosulfan, at the sub-cytotoxic concentration used throughout this study, did not alter cell cycle progression or ciliation, irrespective of whether it was applied before or after ciliogenesis. The reduction in ciliary length, together with the unchanged ciliation and cell cycle data, indicates that endosulfan exerts a direct effect on ciliary integrity rather than producing non-specific cytotoxic effects, providing cellular context for the hedgehog signalling perturbations described below [5].

Across multiple cell lines and tissues, endosulfan reduced GLI1 expression at both transcriptional and protein levels, an effect consistent across qRT-PCR, GLI-responsive luciferase reporter assays, and RNA sequencing of gonadal tissues from exposed mice. Because this suppression occurred independently of ciliation status and could not be rescued by SAG co-treatment, indicating that endosulfan acts downstream of, or in parallel to, SMO activation, we next systematically tested candidate components of the hedgehog pathway to localise its site of action. Molecular docking and MD simulations showed no stable binding of endosulfan to PTCH1, SMO, GPR161, or KIF7, and endosulfan did not affect SAG-induced ciliary accumulation of SMO or ciliary exit of GPR161, indicating that the core ciliary receptor and transducer machinery remains functionally intact. Endosulfan transiently occupied a pocket adjacent to the GLI zinc-finger DNA-binding domain. Still, it exited within the simulated timeframe, consistent with previous reports that endosulfan does not interact directly with DNA [26], and did not affect cAMP levels. These results systematically exclude direct receptor antagonism, GLI–DNA disruption, and cAMP modulation as mechanisms of endosulfan action, thereby localising its effect to the kinases that govern GLI processing.

PKA and GSK3β are constitutively active under basal conditions, PKA through cAMP-dependent dissociation of its catalytic and regulatory subunits, and GSK3β through constitutive Tyr216 autophosphorylation [10,41]. MD simulations showed that endosulfan forms stable, persistent interactions with both kinases, and in vitro kinase assays confirmed that it increases the catalytic activity of both PKA and GSK3β. To our knowledge, no small-molecule activator of GSK3β has been reported, making this a pharmacologically unprecedented and mechanistically significant finding: endosulfan acts as a direct activator of an already active kinase, rather than as a conventional inhibitor or an allosteric activator of a quiescent enzyme.

For PKA-C, endosulfan binding left the global free energy landscape and conformational entropy essentially unchanged, indicating no large-scale conformational transition. RMSF analysis instead revealed a significant, coordinated reduction in flexibility at two functionally coupled catalytic elements, the G-rich loop and the activation loop, a signature consistent with the ordered DFG-in conformation required for full catalytic competence [42,43].

Closure of the G-rich loop over the ATP γ-phosphate is known to propagate an allosteric signal through the catalytic spine to the activation loop [44,45]; the coordinated ordering of both elements observed here suggests that endosulfan stabilises a catalytically optimal, ternary-complex-like geometry, fine-tuning an already-active enzyme rather than switching it into a new state.

GSK3β engaged endosulfan through a distinct mode. The free energy landscape showed that endosulfan binding shifts the conformational equilibrium toward a low-RMSD, active-like basin, which is 100–125 kJ/mol more favourable than the high-RMSD inactive basin preferred by the apo enzyme. This shift indicates that endosulfan acts as a potent conformational selector rather than a weak modulator. This shift occurred without a change in solvent accessibility or global conformational entropy, arguing against an entropy-driven mechanism and instead pointing to enthalpic loss of stabilising contacts present in the inactive ground state as the likely basis for the reduced thermal stability observed by CETSA. A comparable phenomenon has been described for the activating EGFR L858R mutation, which similarly reduces thermal stability by locking the kinase into an active conformation lacking the stabilising contacts of the inactive ground state [46]. This structural prediction was functionally validated: GSK3β activity was significantly increased in the presence of endosulfan, and partial rescue of GLI3 ciliary tip accumulation and GLI1 levels by the GSK3β inhibitor BRD3731 confirmed that endosulfan suppresses hedgehog signalling, at least in part, through GSK3β hyperactivation.

Critically, the kinase-activating effect of endosulfan is not a general property of the compound across protein kinases. Endosulfan did not affect the kinase activity of the CDK1–Cyclin A2 complex in vitro, and 500 ns MD simulations demonstrated that endosulfan failed to form stable complexes with either CDK1 or CDK2, in marked contrast to the stable, persistent GSK3β–endosulfan interaction observed over 500 ns. This kinase selectivity is further supported by DeepKinomeWeb analysis of a 225-kinase panel, which revealed that endosulfan’s predicted binding profile is qualitatively distinct from those of all five reference kinase inhibitors examined and does not show generic active-site engagement across the kinome **(Figure S9)**. The structural basis for this selectivity likely reflects the specific geometry of the endosulfan binding site in GSK3β, which does not overlap with the ATP-competitive binding site of staurosporine **(Figure 6D)**, and the distinct conformational landscape of the GSK3β activation loop compared with CDK1 and CDK2. Together, these analyses show that endosulfan engages PKA and GSK3β through mechanistically distinct strategies: dynamic fine-tuning of an already active kinase in the case of PKA-C, and conformational selection of an active state in the case of GSK3β, that converge on the same functional outcome: hyperphosphorylation-driven suppression of GLI activity. Notably, although endosulfan binds in proximity to the ATP-binding site of both PKA-C and GSK3β, it does not occlude ATP binding, consistent with the increased rather than decreased kinase activity observed *in vitro*. As BRD3731 only partially rescued the endosulfan phenotype, PKA and GSK3β likely act non-redundantly downstream of endosulfan exposure; whether combined inhibition of both kinases achieves full pathway rescue is an important question for future work.

Because GSK3β is a central node in Wnt, insulin, and neurotrophin signalling, with dysregulation implicated in Alzheimer’s disease, type 2 diabetes, bipolar disorder, and multiple cancers, identifying the first small-molecule GSK3β activator has implications that extend well beyond the hedgehog pathway. Therapeutic efforts have focused almost exclusively on GSK3β inhibition; activating compounds have not been reported previously, in part because the kinase is constitutively active under basal conditions. The conformational selection mechanism identified here, whereby a small molecule stabilises the active kinase conformation without occluding ATP binding, represents a pharmacologically distinct mode of kinase modulation. These findings raise the question of whether endogenous regulators exploit the same conformational-selection mechanism, and whether comparable dynamics drive pathological GSK3β hyperactivation in disease, questions with broad relevance to kinase biology and, potentially, to therapeutic design.

Centrosomal proteins are essential for ciliary biogenesis, as the mother centriole gives rise to the cilium upon exit from the cell cycle into G0/G1 [47]. Using convergent consensus motif analysis and GLI1 ChIP-seq data, we identified *Cetn3* as a direct GLI1 target gene. Centrin3 localises to the distal end and central core of the centriole, where it regulates RNA splicing of spindle-, centriole-, and ciliogenesis-related genes, including *Pcm1* and *Cep250* [36,48]. We found that endosulfan reduced Centrin3 expression and disrupted its localisation at the ciliary base. CEP250, which mediates centriole cohesion and whose loss increases centriole-to-centriole distance and causes germ cell depletion and infertility in mice [37,38,49], was also significantly reduced in expression and ciliary base localisation upon endosulfan exposure.

The increased centriolar distance observed in endosulfan-treated IMCD3 and TM3 cells provides a direct mechanistic link between endosulfan exposure, centriole cohesion defects, and the reproductive phenotypes reported in exposed populations, though whether this molecular mechanism causally produces the in vivo reproductive phenotypes remains to be validated genetically in animal models.

Taken together, these findings converge on a coherent mechanistic explanation for aspects of the developmental and reproductive phenotype associated with endosulfan exposure. Kinase-driven suppression of GLI activity and downstream repression of the centrosomal genes *Cetn3* and *Cep250* collectively disrupt centriole cohesion and ciliary integrity in reproductive tissue, providing a candidate molecular basis for the altered sperm morphology, reduced sperm counts, and infertility reported in both endosulfan-exposed human populations and animal models [24–27]. More broadly, this multi-pronged disruption of a signalling axis, at the levels of kinase activity and transcriptional output, illustrates how a single environmental toxicant can perturb developmental signalling through several converging molecular routes. Whether other environmental chemicals similarly target the hedgehog kinase machinery to produce ciliopathy-like molecular phenotypes is an important question raised by these findings.

Several limitations of this study warrant consideration. The kinase selectivity of endosulfan has been assessed using DeepKinomeWeb predictions across 225 kinases, MD simulations against CDK1, CDK2, and SRC **(Movie EV7)**, and in vitro kinase assay validation against GSK3β and CDK1–Cyclin A2. While these analyses collectively support selective GSK3β engagement, comprehensive experimental kinome profiling using biochemical or chemoproteomics approaches, such as KINOMEscan or thermal proteome profiling, would further define the selectivity landscape in a cellular context. GSK3β inhibition by BRD3731 only partially rescued endosulfan-induced GLI suppression, indicating that PKA and GSK3β act non-redundantly; whether combined inhibition of both kinases is sufficient to rescue the pathway fully remains to be tested. Finally, our mechanistic data are derived primarily from cultured cell lines and short-term in vivo exposure in mice; establishing whether the hedgehog pathway suppression observed here causally contributes to the developmental and reproductive phenotypes of endosulfan exposure will require genetic rescue experiments in appropriate animal models.

In summary, this study provides the first mechanistic evidence that an environmental toxicant perturbs primary cilium-mediated hedgehog signalling through selective activation of GSK3β, the first reported small-molecule activator of this kinase, and allosteric fine-tuning of PKA, suppressing GLI activity and the expression of newly identified ciliary and centrosomal GLI targets *Cetn3* and *Cep250* that are required for centriole cohesion (**Figure 7L)**. The kinase selectivity of endosulfan for GSK3β over structurally related kinases, including CDK1 and CDK2, established by both in vitro kinase assays and MD simulations, suggests a specific mode of molecular engagement rather than a generic kinase perturbation. By disrupting this signalling–ciliogenesis feedback loop at multiple convergent points, endosulfan exposure produces molecular phenotypes overlapping with those of genetic ciliopathies, providing a candidate mechanistic framework for understanding pesticide-associated developmental and reproductive toxicity. More broadly, these findings identify the hedgehog kinase machinery as a previously unrecognised target for environmental chemical disruption and suggest that kinase-level interrogation can more generally reveal mechanisms of environmental toxicity missed by conventional receptor-binding or genotoxicity screens.

## Methods

### Reagents and tools table

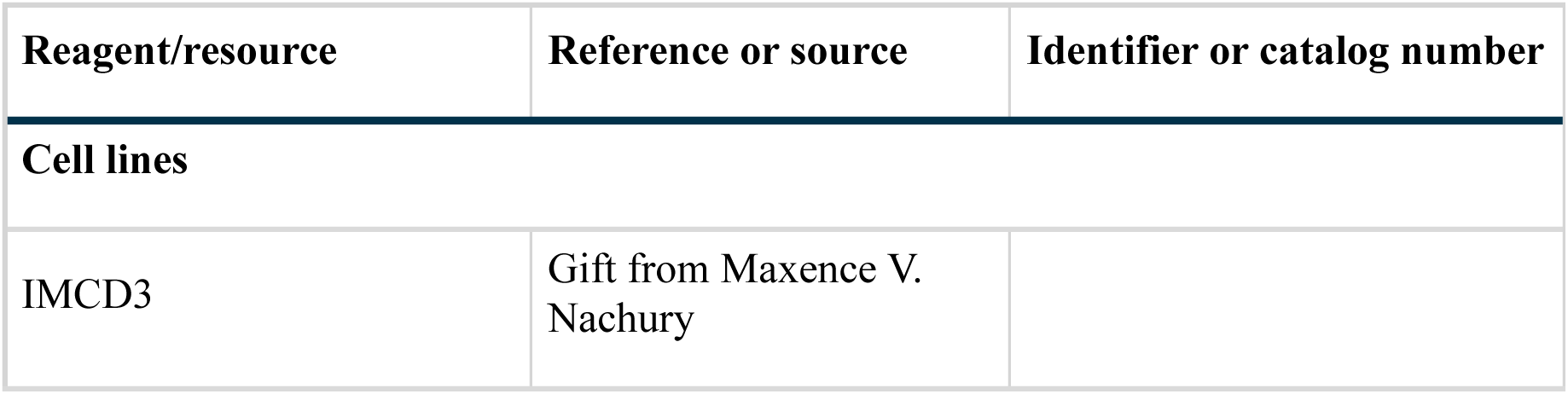
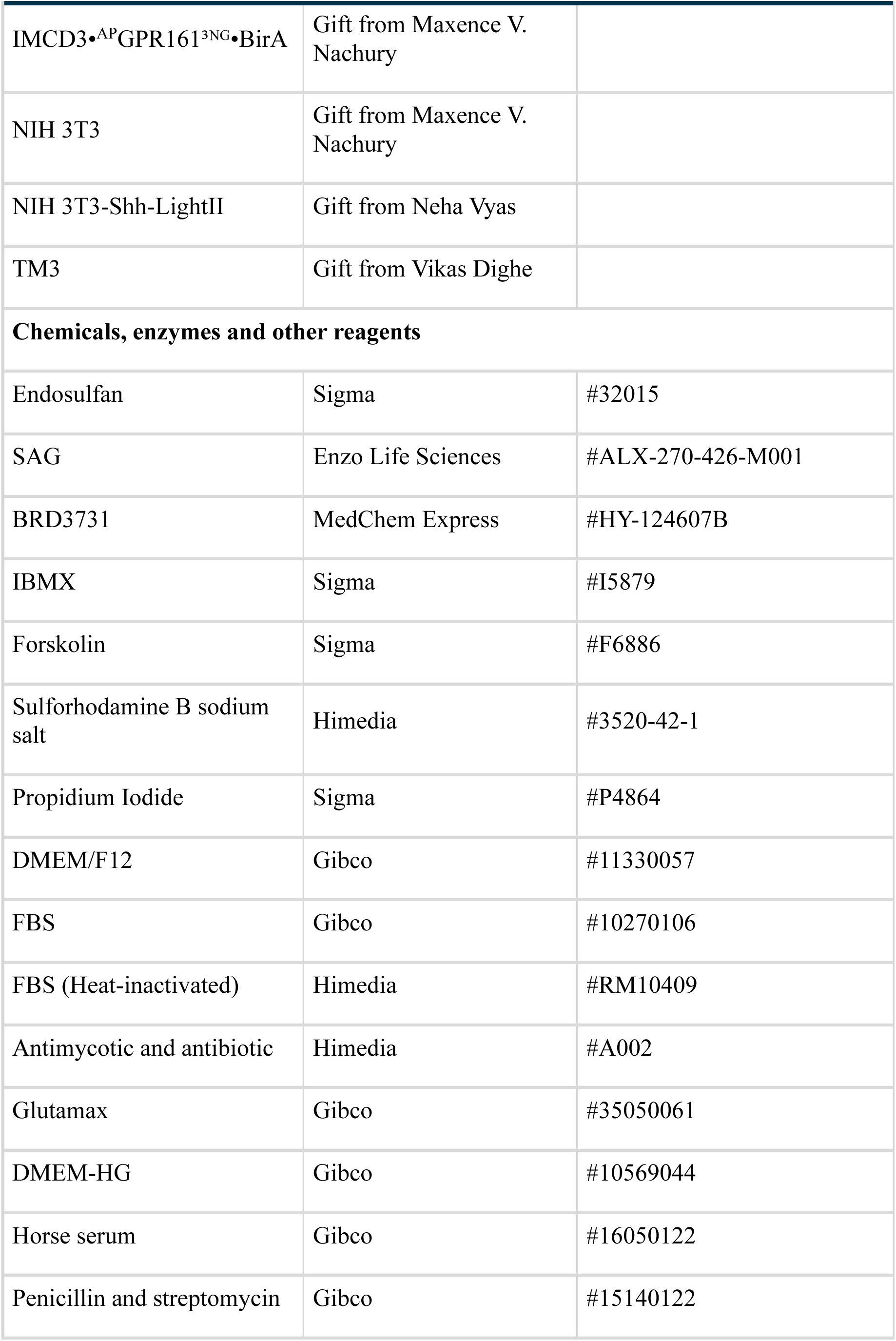
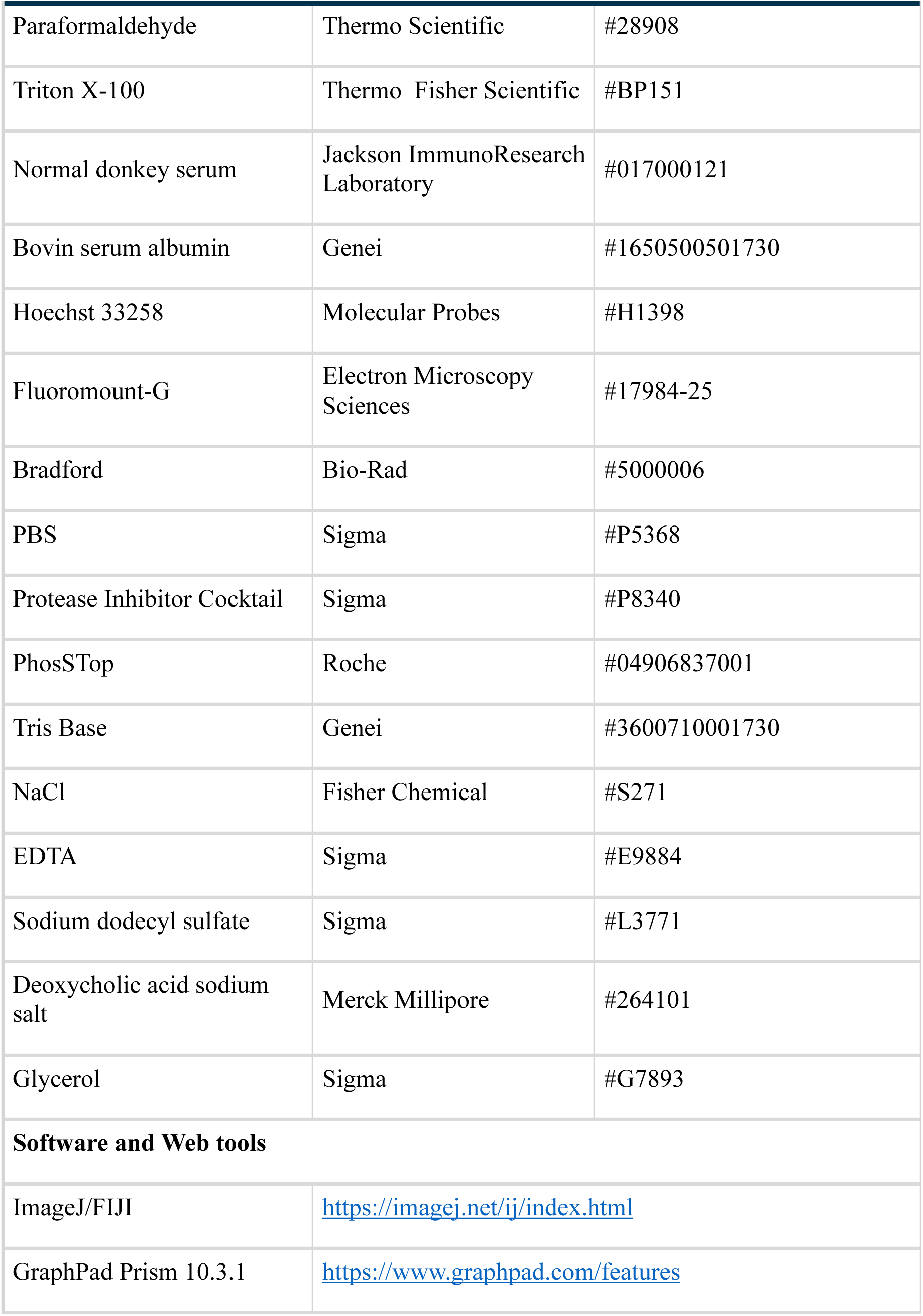
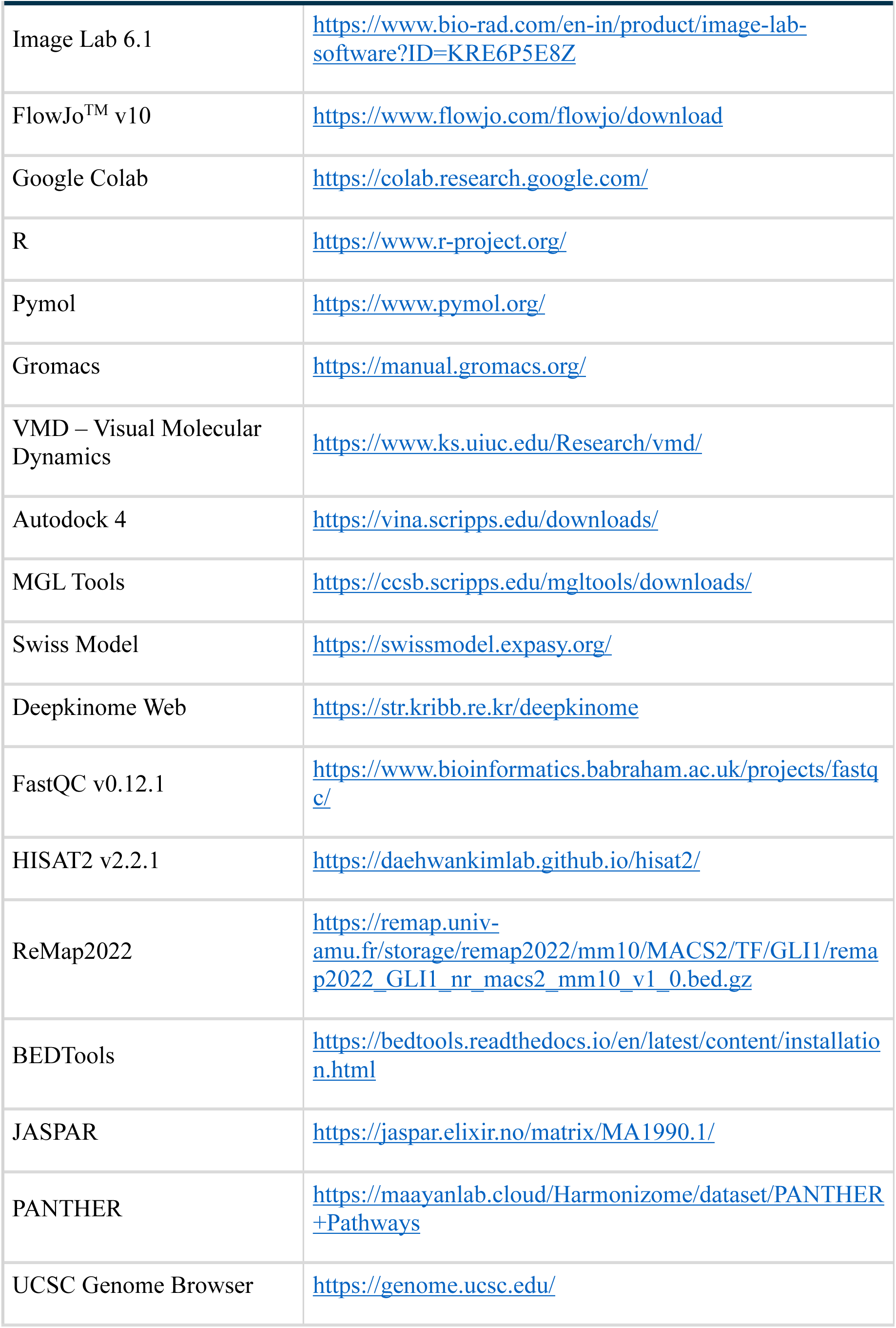
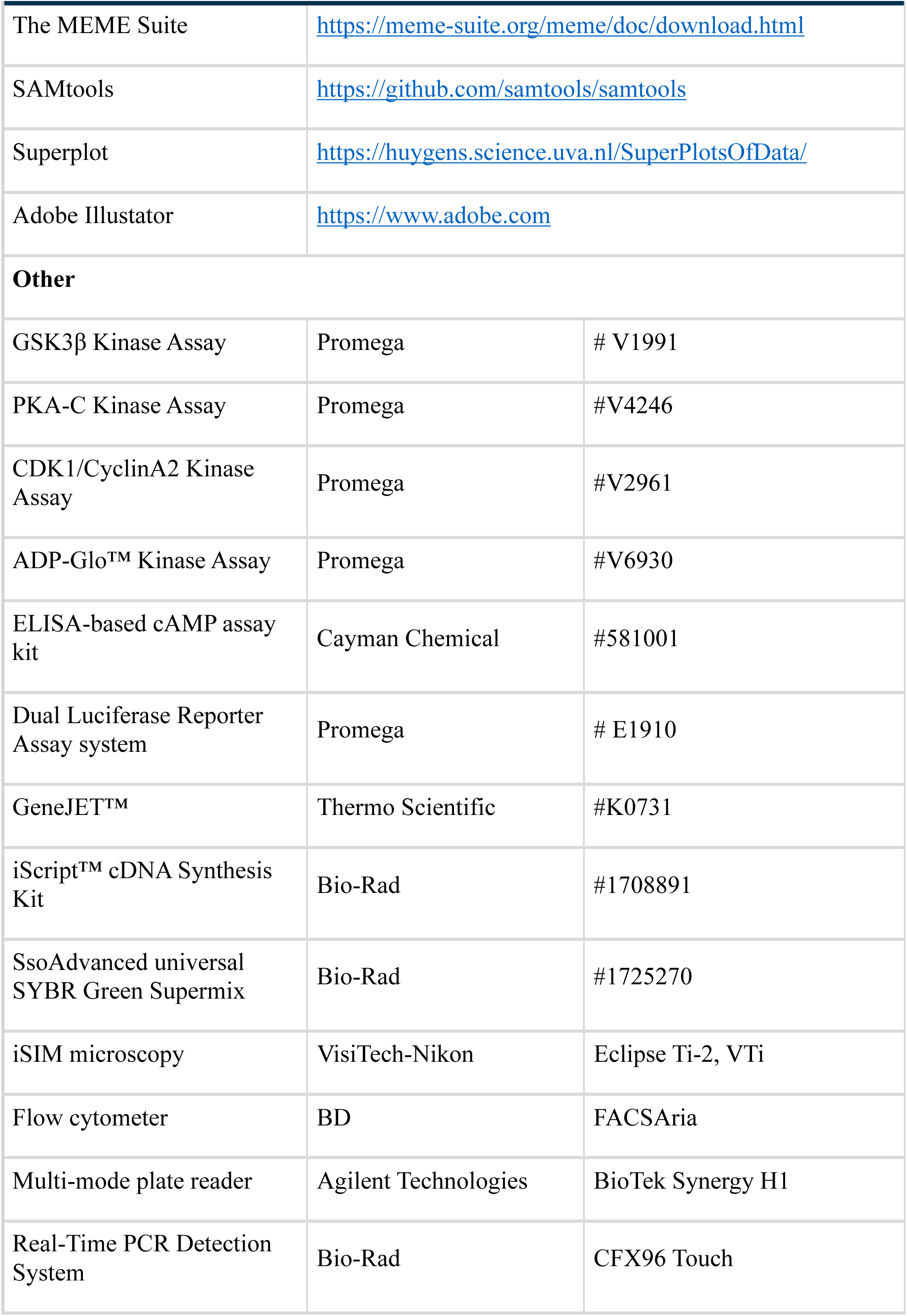
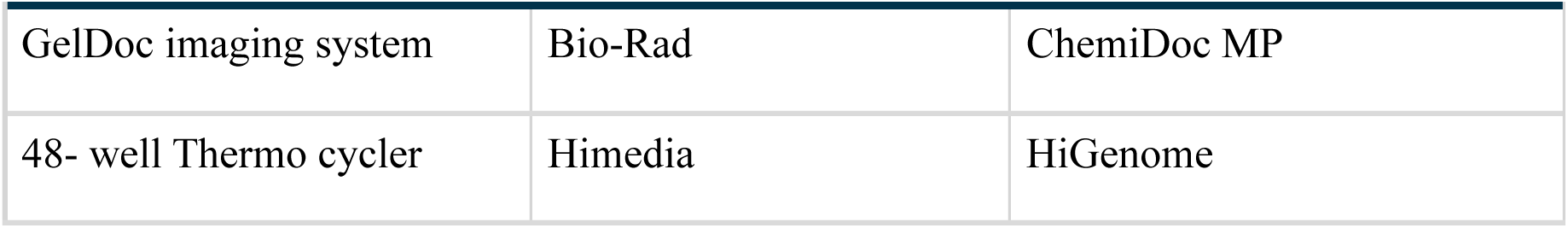

Reagents and tools table:
| Reagent/resource | Reference or source | Identifier or catalog number |
| --- | --- | --- |
| **Cell lines** |  |  |
| IMCD3 | Gift from Maxence V. Nachury |  |
| IMCD3•<sup>AP</sup>GPR161<sup>3NG</sup>•BirA | Gift from Maxence V. Nachury |  |
| NIH 3T3 | Gift from Maxence V. Nachury |  |
| NIH 3T3-Shh-LightII | Gift from Neha Vyas |  |
| TM3 | Gift from Vikas Dighe |  |
| **Chemicals, enzymes and other reagents** |  |  |
| Endosulfan | Sigma | #32015 |
| SAG | Enzo Life Sciences | #ALX-270-426-M001 |
| BRD3731 | MedChem Express | #HY-124607B |
| IBMX | Sigma | #I5879 |
| Forskolin | Sigma | #F6886 |
| Sulforhodamine B sodium salt | Himedia | #3520-42-1 |
| Propidium Iodide | Sigma | #P4864 |
| DMEM/F12 | Gibco | #11330057 |
| FBS | Gibco | #10270106 |
| FBS (Heat-inactivated) | Himedia | #RM10409 |
| Antimycotic and antibiotic | Himedia | #A002 |
| Glutamax | Gibco | #35050061 |
| DMEM-HG | Gibco | #10569044 |
| Horse serum | Gibco | #16050122 |
| Penicillin and streptomycin | Gibco | #15140122 |
| Paraformaldehyde | Thermo Scientific | #28908 |
| Triton X-100 | Thermo Fisher Scientific | #BP151 |
| Normal donkey serum | Jackson ImmunoResearch Laboratory | #017000121 |
| Bovin serum albumin | Genei | #1650500501730 |
| Hoechst 33258 | Molecular Probes | #H1398 |
| Fluoromount-G | Electron Microscopy Sciences | #17984-25 |
| Bradford | Bio-Rad | #5000006 |
| PBS | Sigma | #P5368 |
| Protease Inhibitor Cocktail | Sigma | #P8340 |
| PhosSTop | Roche | #04906837001 |
| Tris Base | Genei | #3600710001730 |
| NaCl | Fisher Chemical | #S271 |
| EDTA | Sigma | #E9884 |
| Sodium dodecyl sulfate | Sigma | #L3771 |
| Deoxycholic acid sodium salt | Merck Millipore | #264101 |
| Glycerol | Sigma | #G7893 |
| **Software and Web tools** |  |  |
| ImageJ/FIJI | https://imagej.net/ij/index.html |  |
| GraphPad Prism 10.3.1 | https://www.graphpad.com/features |  |
| Image Lab 6.1 | https://www.bio-rad.com/en-in/product/image-lab-software?ID=KRE6P5E8Z |  |
| FlowJo™ v10 | https://www.flowjo.com/flowjo/download |  |
| Google Colab | https://colab.research.google.com/ |  |
| R | https://www.r-project.org/ |  |
| Pymol | https://www.pymol.org/ |  |
| Gromacs | https://manual.gromacs.org/ |  |
| VMD – Visual Molecular Dynamics | https://www.ks.uiuc.edu/Research/vmd/ |  |
| Autodock 4 | https://vina.scripps.edu/downloads/ |  |
| MGL Tools | https://ccsb.scripps.edu/mgltools/downloads/ |  |
| Swiss Model | https://swissmodel.expasy.org/ |  |
| Deepkinome Web | https://str.kribb.re.kr/deepkinome |  |
| FastQC v0.12.1 | https://www.bioinformatics.babraham.ac.uk/projects/fastqc/ |  |
| HISAT2 v2.2.1 | https://daehwankimlab.github.io/hisat2/ |  |
| ReMap2022 | https://remap.univ-amu.fr/storage/remap2022/mm10/MACS2/TF/GLI1/remap2022_GLI1_nr_macs2_mm10_v1_0.bed.gz |  |
| BEDTools | https://bedtools.readthedocs.io/en/latest/content/installation.html |  |
| JASPAR | https://jaspar.elixir.no/matrix/MA1990.1/ |  |
| PANTHER | https://maayanlab.cloud/Harmonizome/dataset/PANTHER+Pathways |  |
| UCSC Genome Browser | https://genome.ucsc.edu/ |  |
| The MEME Suite | https://meme-suite.org/meme/doc/download.html |  |
| SAMtools | https://github.com/samtools/samtools |  |
| Superplot | https://huygens.science.uva.nl/SuperPlotsOfData/ |  |
| Adobe Illustator | https://www.adobe.com |  |
| **Other** |  |  |
| GSK3β Kinase Assay | Promega | # V1991 |
| PKA-C Kinase Assay | Promega | #V4246 |
| CDK1/CyclinA2 Kinase Assay | Promega | #V2961 |
| ADP-Glo™ Kinase Assay | Promega | #V6930 |
| ELISA-based cAMP assay kit | Cayman Chemical | #581001 |
| Dual Luciferase Reporter Assay system | Promega | # E1910 |
| GeneJET™ | Thermo Scientific | #K0731 |
| iScript™ cDNA Synthesis Kit | Bio-Rad | #1708891 |
| SsoAdvanced universal SYBR Green Supermix | Bio-Rad | #1725270 |
| iSIM microscopy | VisiTech-Nikon | Eclipse Ti-2, VTi |
| Flow cytometer | BD | FACSAria |
| Multi-mode plate reader | Agilent Technologies | BioTek Synergy H1 |
| Real-Time PCR Detection System | Bio-Rad | CFX96 Touch |
| GelDoc imaging system | Bio-Rad | ChemiDoc MP |
| 48- well Thermo cycler | Himedia | HiGenome |

### Antibodies used in this study

**Antibodies used in this study:**
| Antibody | Species | Dilution | Company | Cat. No. | Application |
| --- | --- | --- | --- | --- | --- |
| Monoclonal Primary |  |  |  |  |  |
| $\alpha$ - tubulin | Mouse | 1:1000 | Sigma | T6199 | WB |
| Acetylated tubulin | Mouse | 1:10000 | Sigma | T7451 | IF |
| γ-tubulin | Mouse | 1:5000 | Sigma | T6557 | IF |
| GLI1 | Mouse | 1:1000 | Cell Signaling Technology | A2643S | WB |
| SMO | Mouse | 1:250 | Santa Cruz Biotechnology | Sc-166685 | IF |
| GSK3β | Rabbit | 1:1000 | Cell Signaling Technology | 12456S | WB |
| β-actin | Mouse | 1:1000 | Santa Cruz Biotechnology | Sc-47778 | WB |
| Polyclonal Primary |  |  |  |  |  |
| IFT88 | Rabbit | 1:200 | Proteintech | 13967-1-AP | IF |
| GLI3 | Goat | 1 μg/mL and 4 μg/mL | R&D Systems | AF3690 | WB and IF |
| γ-tubulin | Rabbit | 1:500 | Abcam | AB11317 | IF and |
| CEP250 | Rabbit | 1:200 | Proteintech | 14498-1-AP | IF |
| ARL13B | Rabbit | 1:1000 | Proteintech | 17711-1-AP | IF |
| Centrin3 | Mouse | 1:500 | Gift from Shubhra Mazumdar |  | IF |
| Fluorophore-tagged secondary |  |  |  |  |  |
| Cy5 | Goat | 1:500 | Jackson ImmunoResearch Laboratory | 115-175-205 | IF |
| Cy5 | Goat |  | Jackson ImmunoResearch Laboratory | 115-175-206 | IF |
| Cy5 | Donkey |  | Jackson ImmunoResearch Laboratory | 711-175-152 | IF |
| Cy3 | Goat |  | Jackson ImmunoResearch Laboratory | 115-165-207 | IF |
| Cy3 | Goat |  | Jackson ImmunoResearch Laboratory | 115-165-206 | IF |
| Cy3 | Donkey |  | Jackson ImmunoResearch Laboratory | 711-165-152 | IF |
| Alexa 488 | Goat |  | Invitrogen | A11001 | IF |
| Alexa 488 | Goat |  | Invitrogen | A11008 | IF |
| Alexa 488 | Donkey |  | Invitrogen | A21202 | IF |
| Alexa 555 | Donkey |  | Invitrogen | A31572 | IF |
| Alexa 633 | Donkey |  | Invitrogen | A21082 | IF |
| HRP-tagged secondary |  |  |  |  |  |
| Anti-mouse | Goat | 1:10000 | Cell Signaling Technology | 7076S | WB |
| Anti-rabbit | Goat |  | Cell Signaling Technology | 7074S | WB |
| Anti-goat | Donkey |  | Jackson ImmunoResearch Laboratory | 705-035-147 | WB |

### Oligos used in this study

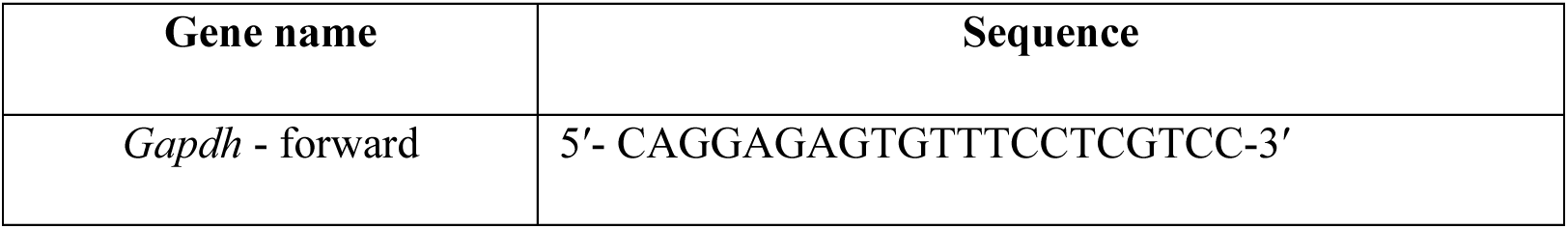
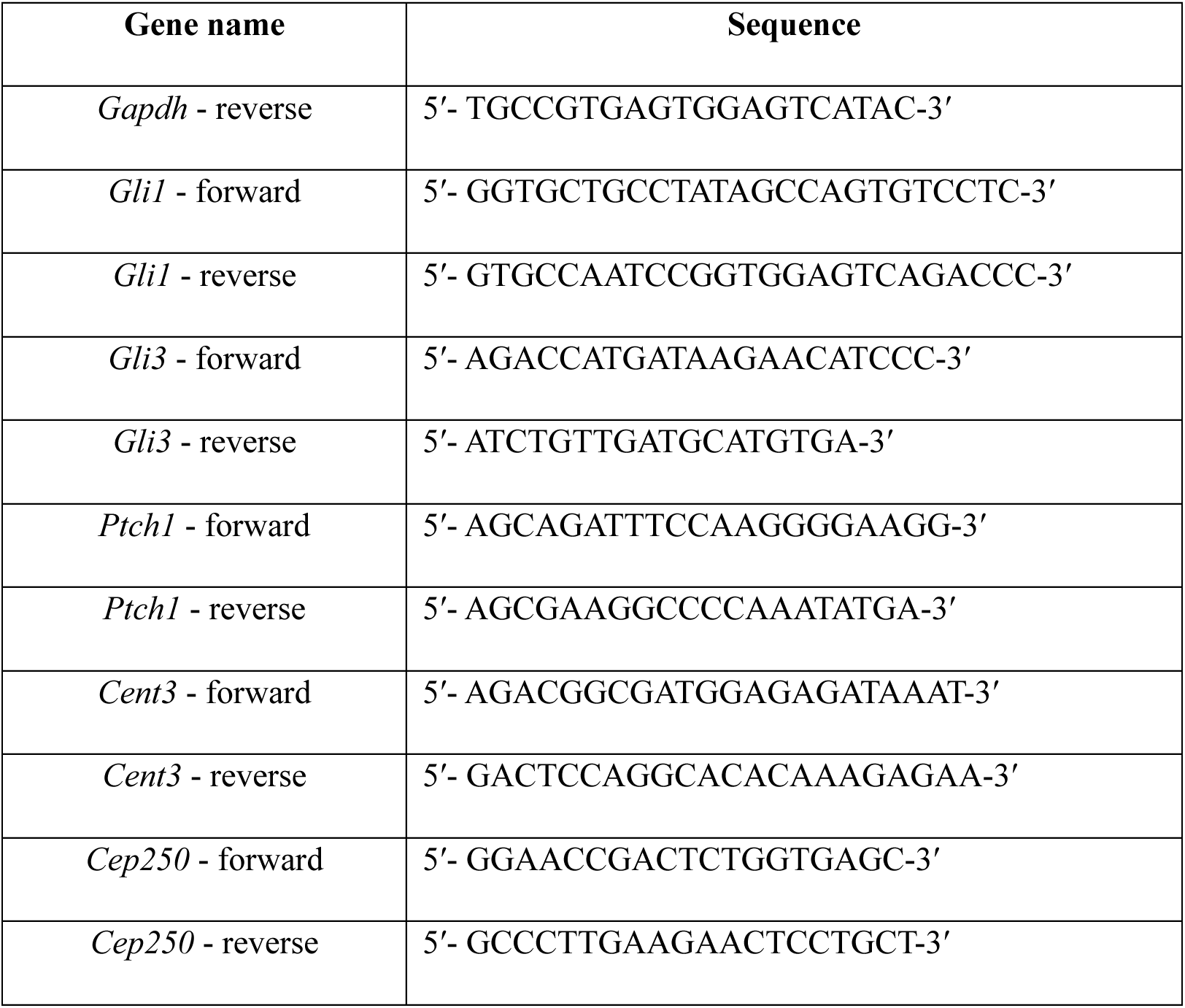

### Cell culture

The parental mouse IMCD3-FlpIn cell line (gift from P.K. Jackson, Stanford University, Stanford, CA) and IMCD3•^AP^GPR161³ᴺᴳ•BirA (gift from Maxence V. Nachury, University of California, San Francisco, CA) cells were cultured in DMEM/F12 (11330057; Gibco) supplemented with 10% FBS (10270106; Gibco), 1x antimycotic and antibiotic (A002; Himedia), and Glutamax (35050061; Gibco). MCF7 cells (gift from Sushil Kumar, Indian Institute of Technology Bombay, Mumbai, India) and NIH3T3 and NIH3T3-Shh-LightII cells (gift from Neha Vyas, St John’s Research Institute, Bangalore, India) were cultured in DMEM-HG (10569044; Gibco) + Glutamax supplemented with 10% FBS and 1x antimycotic and antibiotic. TM3 cells (gift from Vikas Dighe, ICMR National Institute for Research on Women’s Health, Mumbai, India) were cultured in DMEM/F12 supplemented with 5% horse serum (16050122; Gibco), 2.5% FBS, Glutamax and 1x Penicillin and Streptomycin (15140122; Gibco). Ciliation was induced by serum starvation in media containing 0.2% FBS for 16 h. Cells were incubated at 37 °C with 5% CO2.

### Calculation of half-maximal inhibitory concentration

IMCD3•^AP^GPR161³ᴺᴳ•BirA, NIH3T3, and TM3 cells were seeded in 96-well plates at a density of 5,000 cells/well. The half-maximal inhibitory concentration (IC₅₀) of endosulfan (Cat. No. 32015; Millipore-Sigma) was determined using the sulforhodamine B (SRB) assay as described previously [50]. Two treatment conditions were employed: (1) cells were treated with endosulfan for 48 h in complete growth medium; and (2) cells were first serum-starved for 16 h, followed by endosulfan treatment for 8 h in serum-free medium. Each condition was performed in at least three independent experiments.

### Cell cycle analysis

IMCD3-FlpIn cells were treated with either the vehicle control (0.05% DMSO) or 12.5 µM endosulfan for 48 h. Subsequently, cells were trypsinised, washed, and fixed in ice-cold 70% ethanol in 1× PBS supplemented with 0.1% FBS, and stored at −20 °C overnight. Cells were then washed twice with 1x PBS and incubated with 50 µg/mL propidium iodide (Cat. No. P4864) and 1 µg/mL RNase A for 15 min at room temperature before acquisition. Cell cycle analysis was performed using a BD FACSAria flow cytometer (BD Biosciences, San Jose, CA, USA), and data were analysed using FlowJo software (BD Biosciences).

### Antibodies and Drugs

The following primary antibodies were used for immunofluorescence: Monoclonal: anti-acetylated tubulin (mouse; clone 6-11B-1; Cat. No. T7451; Sigma-Aldrich; 1:10,000), anti-SMO (mouse; Cat. No. sc-166685; Santa Cruz Biotechnology; 1:250), anti-γ-tubulin (mouse; Cat. No. T6557; Sigma-Aldrich; 1:5,000), anti-GLI1 (mouse; Cat. No. A2346S; Cell Signaling Technology; 1:1,000). Polyclonal: anti-γ-tubulin (rabbit; Cat. No. AB11317; Abcam; 1:500, gift from Sreelaja Nair, Indian Institute of Technology Bombay, Mumbai, India), anti-IFT88 (rabbit; Cat. No. 13967-1-AP; Proteintech; 1:200), anti-CEP250 (rabbit; Cat. No. 14498-1-AP; Proteintech; 1:200), anti-ARL13B (rabbit; Cat. No. 17711-1-AP; Proteintech; 1:1,000), anti-Centrin-3 (mouse; Santa Cruz Biotechnology; 1:500; gift from Shubhra Mazumdar, Presidency University, Kolkata, India), anti-GLI3 (goat; Cat. No. AF3690; R&D Systems; 1 µg/mL). The following fluorophore-tagged secondary antibodies from Jackson ImmunoResearch Laboratory were used: Cy5 anti-mouse IgG1 (goat; Cat. No. 115-175-205), Cy3 anti-mouse IgG2b (goat; Cat. No. 115-165-207), Cy5 anti-mouse IgG2a (goat; Cat. No. 115-175-206), Cy3 anti-mouse IgG2a (goat; Cat. No. 115-165-206), Cy5 anti-rabbit IgG (donkey; Cat. No. 711-175-152), and Cy3 anti-rabbit IgG (donkey; Cat. No. 711-165-152). The following fluorophore-tagged secondary antibodies from Invitrogen were used: Alexa 488 anti-mouse (goat; Cat. No. A11001), Alexa 488 anti-rabbit (goat; Cat. No. A11008), Alexa 488 anti-mouse (donkey; Cat. No. A21202), Alexa 555 anti-rabbit (donkey; Cat. No. A31572), Alexa 633 anti-goat (donkey; Cat. No. A21082). All secondary antibodies were used in 1:500 dilutions.

The following reagents were used at the concentrations indicated: endosulfan (12.5 µM and 6.25 µM; Cat. No. 32015; Millipore-Sigma), SAG (200 nM; Cat. No. ALX-270-426-M001; Enzo Life Sciences), BRD3731 (10 µM; Cat. No. HY-124607B; MedChemExpress, kind gift from Anirban Banerjee, Indian Institute of Technology Bombay, Mumbai, India), 3-isobutyl-1-methylxanthine (IBMX) (1 mM; Cat. No. I5879; Sigma-Aldrich), and forskolin (10 µM; Cat. No. F6886; Sigma-Aldrich). All reagents were dissolved in DMSO (Cat. No. 276855; Sigma-Aldrich).

### GLI-Luciferase Assay

NIH 3T3-Shh-LightII cells, the GLI-responsive firefly luciferase reporter cell line with a constitutive Renilla-luciferase reporter, were seeded onto a 24-well plate at a density of 6 × 10^4^ cells per well DMEM/F12 media (Cat. No 10565-042, Gibco) containing 10% FBS (Cat. No. A5256701, Thermo Scientific) at 37 °C with 5% CO2. 24 h post plating, the cells were either treated with DMSO vehicle control or 25 µM endosulfan in culture medium containing 0.2% FBS and cultured for an additional 48 h. Subsequently, the endosulfan- and DMSO-treated cells were treated with 200 nM SAG, or DMSO (vehicle control) for an additional 24 h after which, the cells were subjected to the luciferase assay using the Dual Luciferase Reporter Assay system (Cat. No. E1910, Promega) as per the manufacturer’s protocol. Briefly, post treatment, the cells were washed once with DPBS (Cat. No. 14190-250, Gibco) and lysed with 100 µl of 1X Passive Lysis Buffer by gently rocking the plates for 15 min at room temperature (RT). Then, the lysates were collected in 1.5 ml vials, subjected to 6 cycles of freeze-thaw to ensure complete lysis and then, 20 µl of the cell lysate was mixed with 100 µl of the Luciferase Assay Reagent II (LAR II) was added and the firefly luciferase activity was determined using the SIRIUS Luminometer (Berthold Detection Systems). Subsequently, 100 µl of Stop-and-Glo reagent was added and 10 s later, readings recorded again to determine the Renilla luciferase activity. The readings were then used to determine the overall activity of Hedgehog based on the ratio of firefly-to-Renilla luciferase, which was plotted as relative luciferase activity.

### *In vitro* Kinase Assay

The effect of endosulfan on the *in vitro* kinase activity of PKA-C (Cat. No. V4246; Promega), GSK3β (Cat. No. V1991; Promega) and CDK1/CyclinA2 (Cat. No. V2961; Promega) was assessed using the ADP-Glo™ Kinase Assay (Cat. No. V6930; Promega) according to the manufacturer’s protocol. Briefly, the assay proceeds in two sequential steps. In the first step, ATP is consumed during the kinase reaction, generating ADP proportional to kinase activity. In the second step, the remaining ADP is converted back to ATP, which is subsequently utilized in a luciferase-based bioluminescence reaction. The resulting luminescence signal was quantified using a microplate reader (BioTek Synergy H1; Agilent Technologies) and is directly proportional to PKA, GSK3β, and CDK1 kinase activity.

### ELISA-based cAMP Assay

IMCD3-FlpIn cells were seeded at a density of 40,000 cells/well in 24-well plates in complete growth medium. Endosulfan treatment was administered under two conditions: (1) endosulfan treatment for 48 h followed by serum starvation for 16 h; and (2) endosulfan treatment for 48 h, serum starvation for 16 h, followed by a second endosulfan treatment for 10 min. Cells were then incubated in stimulation medium, serum-free, antibiotic-free medium supplemented with 0.3% BSA and 1 mM IBMX, for 30 min at 37 °C. To measure basal and maximum cAMP levels, cells were supplemented with stimulation medium alone or stimulation medium containing 10 µM forskolin, respectively. Forskolin was applied under two conditions: (1) a 10-minute treatment in fresh stimulation medium following serum starvation, and (2) a 30-minute treatment in fresh stimulation medium following serum starvation. In both conditions, cells were washed twice with ice-cold PBS following treatment. Intracellular cAMP levels were measured using an ELISA-based cAMP assay kit (Cat. No. 581001; Cayman Chemical) according to the manufacturer’s protocol. Absorbance was recorded at 412 nm using a microplate reader (BioTek Synergy H1; Agilent Technologies). cAMP responses were normalized to basal cAMP levels prior to plotting.

### qRT-PCR

Total RNA was extracted from IMCD3-FlpIn cells following 48 h of endosulfan treatment using the GeneJET™ RNA Purification Kit (Cat. No. K0731; Thermo Fisher Scientific) according to the manufacturer’s instructions. Complementary DNA (cDNA) synthesis was performed using the iScript™ cDNA Synthesis Kit (Cat. No. 1708891; Bio-Rad, Hercules, CA, USA) according to the manufacturer’s protocol. Quantitative real-time PCR (qRT-PCR) was performed using SsoAdvanced universal SYBR Green Supermix (Cat. No. 1725270; Bio-Rad) in a total reaction volume of 10 µL using a CFX96 Touch Real-Time PCR Detection System (Bio-Rad). For each reaction, 30 ng of single-stranded cDNA was used as template. *Gapdh* was used as an internal reference gene for normalization. Relative mRNA expression levels were calculated using the 2^−ΔΔCt^ method. Each sample was analyzed in technical triplicate, and mean values were reported. The primers used in this study are listed in the resource table.

### Immunofluorescence and Imaging

70,000 cells were seeded onto ethanol-sterilized coverslips (12 mm, No. 1.5; Cat. No. 12-545-81; Thermo Fisher Scientific) and allowed to adhere for 12 h in a humidified incubator at 37 °C with 5% CO₂. Under condition 1, cells were treated with either vehicle (0.05% DMSO, v/v) or endosulfan (12.5 µM) for 48 h in complete growth medium, followed by serum starvation for 16 h. Under condition 2, cells were first serum-starved for 16 h and subsequently treated with either vehicle (0.025% DMSO, v/v) or endosulfan (6.25 µM) for 8 h in serum-free medium. Following treatment, cells were fixed with 4% paraformaldehyde (Cat. No. 28908; Thermo Scientific) in PBS for 15 min at 37 °C, then washed twice with PBS. Cells were subsequently extracted in ice-cold methanol (pre-chilled at −20 °C) for 5 min, followed by three washes with PBS. Cells were then blocked and permeabilized in immunofluorescence (IF) buffer; PBS supplemented with 0.1% Triton X-100 (Cat. No. BP151; Thermo Fisher Scientific), 5% normal donkey serum (Cat. No. 017-000-121; Jackson ImmunoResearch Laboratories), and 3% BSA (Cat. No. 1650500501730; Genei), for 30 min, followed by overnight incubation at 4 °C with primary antibodies diluted in IF buffer. Coverslips were then washed three times with IF buffer and incubated with appropriate species-specific secondary antibodies (Jackson ImmunoResearch Laboratories) diluted in IF buffer for 30 min at room temperature, followed by three washes with IF buffer. Nuclear DNA was stained with Hoechst 33258 (Cat. No. H1398; Molecular Probes) for 4 min at room temperature, followed by two washes with PBS. Coverslips were mounted onto glass slides using Fluoromount-G mounting medium (Cat. No. 17984-25; Electron Microscopy Sciences) and allowed to cure before imaging. Cells were imaged on an instant structured illumination microscope (iSIM; VT-iSIM, Visitech International, equipped on a Nikon Ti2, Nikon Instruments, Tokyo, Japan) using a 60× Plan-Apochromat 1.4 NA oil immersion objective. Z-stacks were acquired at 0.5 µm intervals.

### Western Blotting and Densitometry

For GLI3 detection, cell lysates were prepared by resuspending cells in RIPA buffer (50 mM Tris-HCl pH 7.5, 150 mM NaCl, 1 mM EDTA, 1% Triton X-100, 0.1% SDS, and 1% sodium deoxycholate) supplemented with freshly added phosphatase and protease inhibitors, and incubated on ice for 20 min. Insoluble debris was removed by centrifugation at 21,000 × g for 20 min at 4 °C, and the clarified supernatant was retained.

For GLI1 detection, cells were lysed in SDS lysis buffer (4% SDS, 16% glycerol, and 40 mM Tris-HCl pH 6.8) supplemented with freshly added phosphatase and protease inhibitors, and heated at 95 °C for 15 min. Insoluble debris was removed by centrifugation at 21,000 × g for 20 min at 4 °C, and the clarified supernatant was retained.

Protein concentrations were determined using the Bradford assay with BSA as a standard curve prepared in RIPA buffer. Prior to electrophoresis, samples were denatured by boiling at 95 °C for 5 min in SDS sample loading buffer and placed on ice. Equal amounts of total protein (30 µg per lane) were resolved by SDS-PAGE and analyzed by immunoblotting using the following primary antibodies: anti-GLI3 (goat; Cat. No. AF3690; R&D Systems; 1 µg/mL), anti-GLI1 (mouse; clone L42B10; Cat. No. 2643; Cell Signaling Technology; 1:1,000), and anti-β-actin (mouse; clone C4; Cat. No. sc-47778; Santa Cruz Biotechnology; 1:1,000). The following HRP-linked secondary antibodies were used: anti-mouse IgG (Cat. No. 7076S, Cell Signaling Technology), anti-rabbit IgG (Cat. No. 7074S, Cell Signaling Technology), anti-goat IgG (Cat. No. 705-035-147, ImmunoResearch Laboratory). All secondary antibodies were used in 1:10000 dilutions. Blots were developed using a ChemiDoc imaging system (Cytiva) and a ChemiDoc MP imaging system (Bio-Rad).

Western blot analysis and quantification were performed using ImageLab 6.1 (Bio-Rad). The integrated density of bands for proteins of interest was normalised to the integrated intensity of the loading control protein (β-actin). The normalised value was used to compare changes in total protein levels across conditions. Uncropped blots are shown in the **Figure EV12**.

### Cellular Thermal Shift Assay (CETSA)

IMCD3-FlpIn cells were seeded in 10 cm dishes and grown to 80–90% confluency, at which point endosulfan was added for 1 h. Cells were harvested by trypsinization, washed, and resuspended in PBS. Aliquots of 1.5 × 10⁶ cells were transferred to PCR tubes and heated across a temperature gradient ranging from 37 °C to 67 °C in 5 °C increments for 3 min using a 48-well thermocycler. Cells were subsequently allowed to cool to 25 °C for 3 min. Cell lysis was achieved by three cycles of rapid freeze-thaw using liquid nitrogen. Lysates were clarified by centrifugation at 10,000 × g for 20 min at 4 °C, and the supernatants were retained. Protein concentrations were determined using the Bradford assay with BSA as a standard curve prepared in PBS, using the 37 °C and 42 °C samples as concentration references for normalization across all temperature points. Before electrophoresis, samples were denatured by boiling at 95 °C for 5 min in SDS sample loading buffer and placed on ice. Equal amounts of total protein (30 µg per lane) were resolved by SDS-PAGE and analyzed by immunoblotting using an anti-GSK3β antibody (rabbit; Cat. No. 12456S; Cell Signaling Technology; 1:1,000; kind gift from Anirban Banerjee, Indian Institute of Technology Bombay, Mumbai, India). Blots were developed using a ChemiDoc MP imaging system (Bio-Rad).

### Screening of kinase-inhibitor binding affinity

To assess the binding affinity of endosulfan relative to known GSK3β inhibitors, we compared its predicted kinase-binding profile against those of BRD3731 and staurosporine using the DeepKinomeWeb server [51]. DeepKinomeWeb server is a web-based platform that employs a deep learning regression model to perform high-throughput, quantitative prediction of compound–kinase binding affinities across a predefined kinase panel. Briefly, compound structure files in SDF format were uploaded to the platform, which automatically screens each compound against the kinase panel and generates binding affinity predictions. The primary output is visualised as a heatmap displaying predicted binding affinities for each compound across the kinase panel.

### Transcriptome analysis

Raw RNA sequencing (RNA-seq) FASTQ files were obtained from the dataset and assessed for sequence quality using FastQC v0.12.1 [52]. Adapter sequences and low-quality bases were trimmed using Trimmomatic v.039 [53] with default quality filtering parameters. The filtered reads were aligned to the mouse genome (GRCm39/mm39) using a splice-aware aligner, HISAT2 v2.2.1 [54] with default parameters. The resulting alignment files in SAM (Sequence Alignment Map) were converted to BAM format, sorted by genomic coordinates and later indexed using SAMtools v1.17 [55]. Gene-level read quantification was performed using featureCounts (Subread package) [56], with gene annotation obtained from the mm39 GTF file downloaded from the UCSC Genome Browser. Raw read counts were normalised to Reads Per Kilobase of transcript length per Million reads (RPKM) to account for differences in gene length and sequencing depth. Gene sets corresponding to the Hedgehog signalling pathways were downloaded from the PANTHER pathways gene set library collection curated by the Ma’ayan Laboratory [57,58]. The normalised expression matrix was filtered to retain only genes annotated to the pathway for downstream analysis.

All subsequent analyses and visualisation were performed in R v4.4.2 [59]. Expression values were then log2-transformed (i.e., log2(X+1)), and genes exhibiting zero variance across all samples were excluded prior to visualisation. Heatmaps were then generated from the filtered, log-transformed expression matrices using the pheatmap package in R (v1.0.12), with row-wise z-score scaling.

### Gli1 consensus motif and ChIP peak analysis

A curated list of 238 genes encoding proteins localising to various compartments of the cilium, including the axoneme, microtubules, spindle, and motile cilium, was assembled for promoter motif analysis. A promoter region spanning 2,000 nucleotides (−1,500 to +500 nucleotides relative to the transcription start site, TSS) was defined for each gene. MEME Suite-based motif analysis was performed in Python (Google Colab) to identify consensus motifs enriched across these promoter regions, with particular focus on sequences conforming to the GLI transcription factor binding motif.

Mouse GLI1 ChIP-seq peaks were downloaded from the ReMap2022 database [60]. These peaks were intersected with the 2 kb upstream regions of the TSS of a curated list of 99 ciliary genes (including 31 HH genes and 70 DEGs from RNA-seq) using BEDTools (v2.29.1) [61]. Of these, 55 (18HH and 37 DEGs) upstream regions overlapped with GLI1 peaks. Motif scanning of these 55 overlapping regions using FIMO (MEME Suite) [62] identified 14 genes harbouring a GLI1 binding motif in their promoter regions (**Table EV3 and 4**). The URLs for Gli1 peaks and motif are below.

Gli1 peaks URL: https://remap.univ-amu.fr/storage/remap2022/mm10/MACS2/TF/GLI1/remap2022_GLI1_nr_macs2_mm10_v1_0.bed.gz

Gli1 motif: https://jaspar.elixir.no/matrix/MA1990.1/

### MD Simulation

#### All-atom Molecular Dynamics Simulation of Endosulfan-docked GLI/DNA complex

The crystal structure of the five-finger GLI/DNA complex (PDB ID: 2GLI), determined by X-ray crystallography at a resolution of 2.60 Å, was retrieved from the RCSB Protein Data Bank. A system in which endosulfan (S68) was docked between zinc finger domains 2 and 3 (ZNF2 and ZNF3) of GLI was used as the starting configuration to elucidate the effect of endosulfan on GLI–DNA binding. Protein preparation was carried out using the Protein Preparation Wizard implemented in Schrödinger Maestro, which included the addition of hydrogen atoms, assignment of bond orders, and optimization of protonation states at physiological pH. The prepared structure was energy-minimized using the OPLS4 force field. System preparation for molecular dynamics (MD) simulations was performed using the System Builder module in Desmond (Schrödinger Suite, v2025-3). The system was solvated in a cubic periodic box using the SPC water model, with a minimum distance of 10 Å between the solute and each face of the box. System charge neutralization was achieved by adding Mg²⁺ counter-ions, and the ionic strength was set to 0.15 M. The OPLS4 force field was applied to all components of the system. The solvated system was subsequently relaxed using Desmond’s default relaxation protocol before production simulations. Three independent production MD simulations were performed for each system for 400 ns under the NPT ensemble at a temperature of 300 K and a pressure of 1.01325 bar. Trajectory frames were recorded at 100 ps intervals for subsequent analysis. Simulation stability and key molecular interactions were assessed using the Simulation Interaction Diagram (SID) tool in Maestro. Root mean square deviation (RMSD) plots were generated from the trajectory data using a custom R script.

#### All-atom Molecular Dynamics Simulation of Endosulfan-docked Hedgehog components complex

MD simulations were performed for both the apo form and the endosulfan–protein complex for the following systems and durations: KIF7 (PDB ID: 4A14), SRC (PDB ID: 2H8H), and AR (PDB ID: 2PIT) for 100 ns; PKA (PDB ID: 6E99) for 400 ns; GSK3β (PDB ID: 8FF8); CDK1(PDB ID: 6GU2); and CDK2 (PDB ID: 1H1P) for 500 ns. All simulations were carried out using GROMACS (v2019.1) in triplicate except for KIF7 and SRC. Of the 10 docked poses generated for each ligand, the pose with the highest docking score was selected for MD simulation. Ligand topology files were generated using the ACPYPE automated web server [63], and protein topology files were prepared using the CHARMM27 force field [64] implemented in GROMACS. Each system was solvated in a triclinic periodic box and neutralized by the addition of Na⁺ and Cl⁻ ions to a final concentration of 0.15 M. Energy minimization was performed using the steepest descent algorithm with the Verlet cut-off scheme, with convergence set at a maximum force tolerance of 10 kJ mol⁻¹ nm⁻¹. System equilibration was conducted in two sequential phases: NVT (constant number of particles, volume, and temperature) followed by NPT (constant number of particles, pressure, and temperature), each for 100 ps (50,000 steps). Temperature was maintained at 300 K using the V-rescale thermostat (modified Berendsen method) with a coupling time constant of 0.1 ps, and pressure was maintained at 1 bar using the Parrinello–Rahman barostat [65,66]. Trajectory coordinates were saved every 2 ps. Following equilibration, production MD simulations were performed for the durations specified above. Trajectory analyses were conducted using built-in tools in GROMACS (v2019.1) and Visual Molecular Dynamics (VMD). Molecular visualization images were rendered using PyMOL [67]. RMSD, radius of gyration (Rg), and free energy landscape plots were generated using a custom Python script in Google Colab. All remaining graphs were plotted using GraphPad Prism (v10.0). All simulations were performed on the Spacetime High-Performance Computing (HPC) cluster at the Indian Institute of Technology Bombay (IIT Bombay), Mumbai, India.

### Image analysis

Image files were imported into ImageJ/Fiji (National Institutes of Health, Bethesda, MD, USA) for analysis. For quantification of ciliary length and fluorescence signal intensity in fixed cells, maximum intensity projections (MIPs) were used except for GLI3. For GLI3-positive cilia, the best z-plane was used. Ciliary length was measured manually from the base to the tip of each cilium using the line profile tool. Ciliary GPR161 fluorescence intensity (F_GPR161_) was calculated using the following equation:

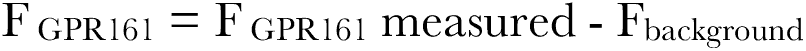

F_GPR161_ is the total ciliary GPR161 fluorescence detected along the cilium, and F_background_ is the GPR161 background fluorescence detected adjacent to the cilium. The fluorescence integrated density was used for all intensity measurements. The same approach was applied to quantify the ciliary fluorescence intensities of IFT88, SMO, Centrin-3, and CEP250. Data were plotted as SuperPlots using the SuperPlotsOfData web application [68]. Each SuperPlot displays the full distribution of individual data points, with the mean value of each independent experiment represented by a larger symbol. All representative micrographs within each panel are displayed using the same dynamic range.

### Statistical analysis

All curve fits for IC_50_ were performed using GraphPad Prism (version 10.3.1). The non-linear regression curve fit model with log (inhibitor) vs. response - Variable slope (four parameters) least squares fit method was chosen. For T_agg,_ the non-linear regression curve-fit model with a Boltzmann sigmoidal form was chosen. We used an unpaired Student’s t-test between two experimental groups, and for more than two conditions, we performed one-way ANOVA followed by Tukey’s post hoc test to determine significance using GraphPad Prism. In bar graphs, data are represented as mean ± SD, and in the superplot, data are represented as mean ± SEM. Asterisks indicate statistical significance values calculated by unpaired Student’s t-test and one-way ANOVA followed by Tukey’s post-hoc multiple comparison test. p-value ≤ 0.0001; ****, p-value ≤ 0.001; ***, p-value ≤ 0.01; **, p-value ≤ 0.05; *.

## Author contributions

R.P. and S.R.S. designed the experiments; R.P. performed all experiments, except those noted below. A.B., A.G., and S.G. conducted the experiments shown in Figure EV3F. S.C.R. and B.C. conducted the experiments shown in Figures 2A and EV4A. H.B. and S.R. conducted the experiments shown in Figures EV5D and EV10. P.B. assisted with the analysis of the computational experiments. R.P. and S.R.S. wrote the manuscript. P.B., S.C.R., B.C., S.G., S.R., and S.R.S. provided technical assistance and experimental resources. R.P. and S.R.S. conceptualised the study. S.R.S. supervised the research and secured funding.

## Acknowledgement

We thank Sathees Raghavan for providing endosulfan; Maxence Nachury for kindly providing the IMCD3-FlpIn and ^AP^GPR161^3NG^ cell lines and the gift of secondary antibodies used for immunofluorescence; Vikash Dighe for kindly providing the TM3 cell line; Anirban Banerjee for the gift of GSK3β antibody and inhibitor; and Sreelaja Nair and Shubhra Mazumdar for providing antibodies. We thank Sudipto Roy, Krishanu Ray, Subba Rao Gangi Shetty, Swadhin Chandra Jana, Radhika Subramanian, Nathalie Jurisch-Yaksi, Chinmay Kamale and the members of Cilia Biology Lab for valuable scientific discussions and critical reading of the manuscript. Imaging was performed on the iSIM microscope at the Research Infrastructure Facility Centre (RIFC), IIT Bombay (IE/19-RIFC0000-CF-04SRFM). Flow cytometry was performed at the FACS Central Facility, IIT Bombay, supported by DST FIST (SR/FST/LSI-055/2000). Computational resources were provided by the Spacetime High-Performance Computing cluster at IIT Bombay. AB acknowledges the University Grants Commission Research Fellowship from the Government of India. SG is supported by iBRIC-inStem core funds, DBT/Wellcome Trust India Alliance Intermediate Fellowship (IA/I/22/1/506238) and the start-up research grant (SRG/2023/000847) from the Science and Engineering Research Board (SERB), Department of Science and Technology, India. RP acknowledges a Senior Research Fellowship from the Department of Biotechnology (DBT), Government of India (DBT/2021-22/IIT-B/1661). This work was supported by the following grants to SRS: a seed grant from IIT Bombay (RD/0522-IRCCSHO-026); a Core Research Grant (CRG/2023/002870) from the Science and Engineering Research Board (SERB), Department of Science and Technology, India; and the Department of Biotechnology (Grant No. BT/PR56541/BMS/85/481/2024). The authors used Grammarly for grammar and spell-checking, and Claude (Anthropic) for assistance with manuscript restructuring. All content was reviewed, verified, and approved by the authors, who take full responsibility for the integrity of the work.

## Expanded View Figures

**Figure EV1:**
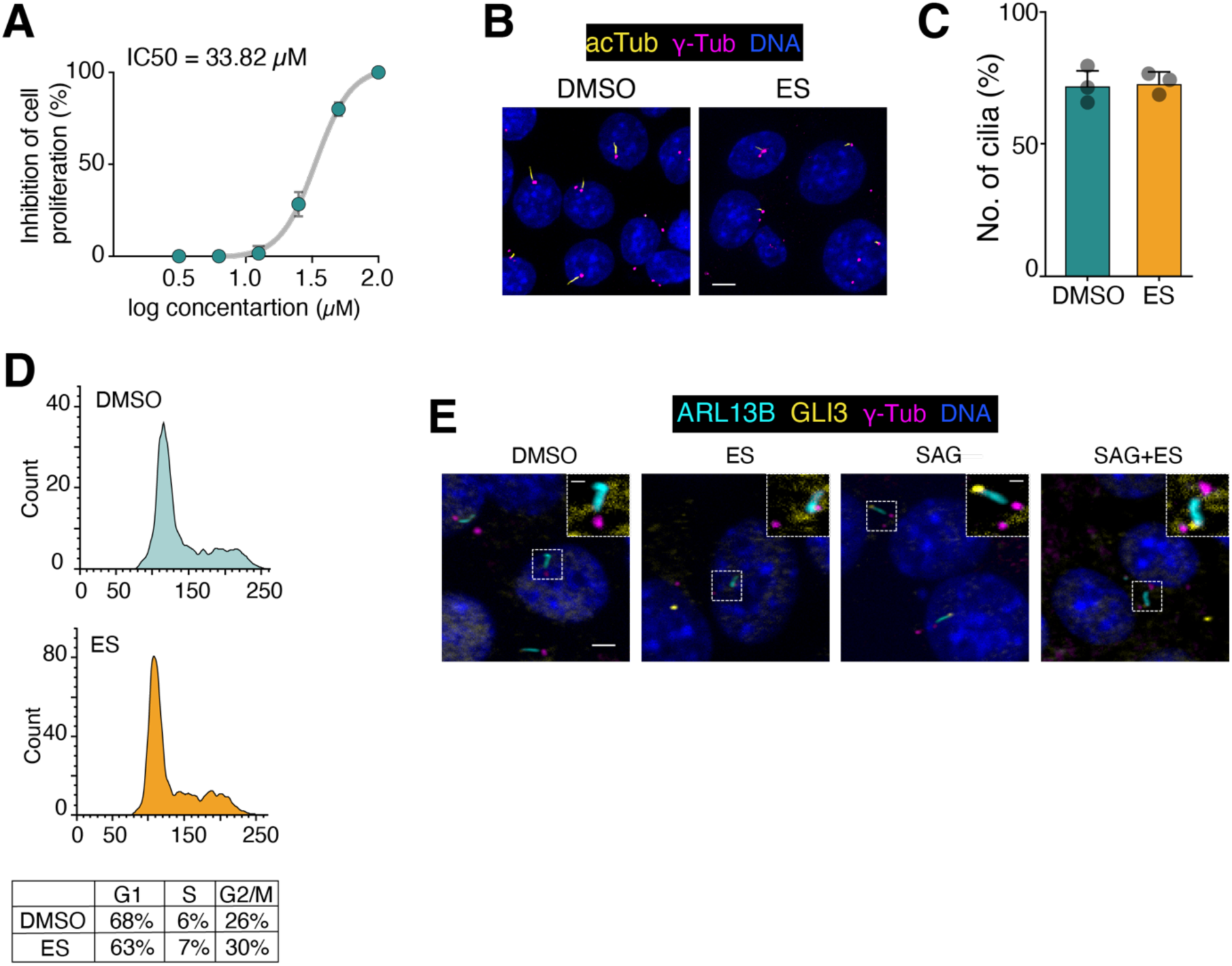
Effect of endosulfan on ciliation and cell cycle progression in IMCD3. **(A)** Cytotoxicity assay showing IC_50_ determination for endosulfan in IMCD3 cells after 48 h of endosulfan treatment. **(B)** Representative immunofluorescence micrographs showing acTub (yellow), γ-tubulin (γ-Tub, magenta), and DNA (Blue). **(C)** Quantification of the percentage of ciliated cells in IMCD3 cells treated with endosulfan or vehicle. Data represent the percentage of ciliated cells from three independent experiments ± SD. **(D)** Cell cycle profile of IMCD3 cells after 48 h of endosulfan treatment. **(E)** A corresponding figure for Figure 1H. Representative immunofluorescence micrographs showing ARL13B (cyan), GLI3 (yellow), and γ-Tub (magenta) in IMCD3 treated as indicated. Scale bars, 5 µm; inset scale bars, 1 µm.

**Figure EV2:**
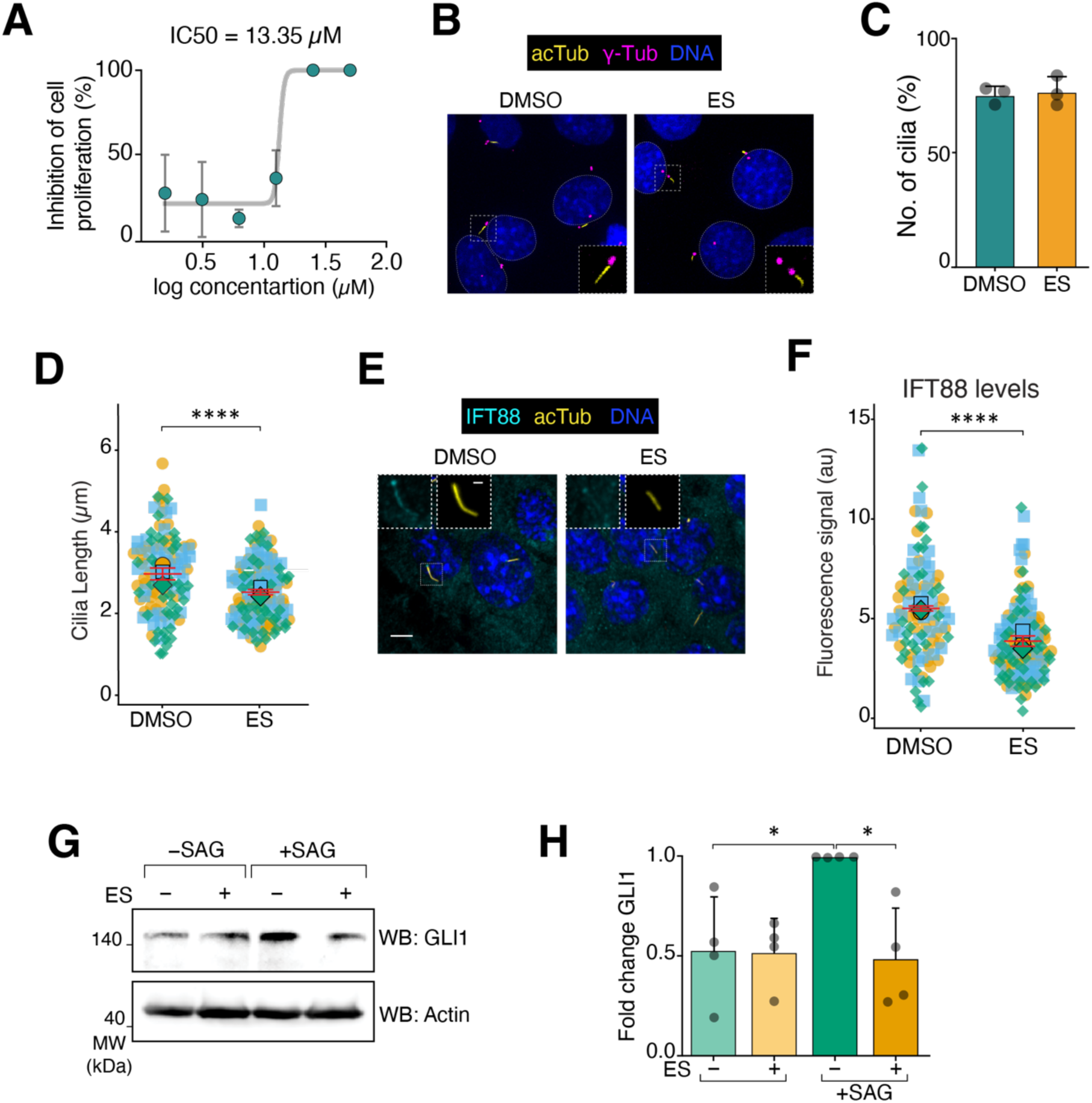
Effect of endosulfan applied post-serum starvation on Hedgehog signalling and cilia. **(A)** Cytotoxicity assay showing IC50 determination for endosulfan in IMCD3 cells after 8 h of endosulfan treatment following 16 h of serum starvation. **(B)** IMCD3 cells treated as indicated were stained for acTub (yellow), γ-Tub (magenta), and DNA (blue). **(C)** Percentage of ciliated cells quantified from (**B**). Data represent the percentage of ciliated cells from three independent experiments, mean ± SD. **(D)** Quantification of ciliary length measured from acTub (panel **B**). Superplots represent all data points from three independent experiments ±SEM. Cell numbers: DMSO, n = 118; endosulfan, n = 133. **(E)** Representative immunofluorescence micrographs showing IFT88 (cyan), acTub (yellow) and DNA (blue). **(F)** Quantification of ciliary IFT88 levels in IMCD3 cells treated with endosulfan or vehicle following serum starvation. Cell numbers: DMSO n = 101, endosulfan n = 136. **G** and **H**) Representative immunoblots **(G)** and quantification of GLI1 protein levels **(H)** from IMCD3 cells treated with endosulfan or vehicle following serum starvation. GLI1 expression in the SAG condition was set to 1. Data represent the mean ±SD of four independent experiments. Fluorescence intensity is expressed in arbitrary units (a.u.). Scale bars, 5 µm; inset scale bars, 1 µm. Asterisks denote statistical significance as determined by unpaired Student’s t-test or one-way ANOVA followed by Tukey’s post-hoc multiple comparison test, as appropriate: **** p ≤ 0.0001; *** p ≤ 0.001; ** p ≤ 0.01; * p ≤ 0.05.

**Figure EV3:**
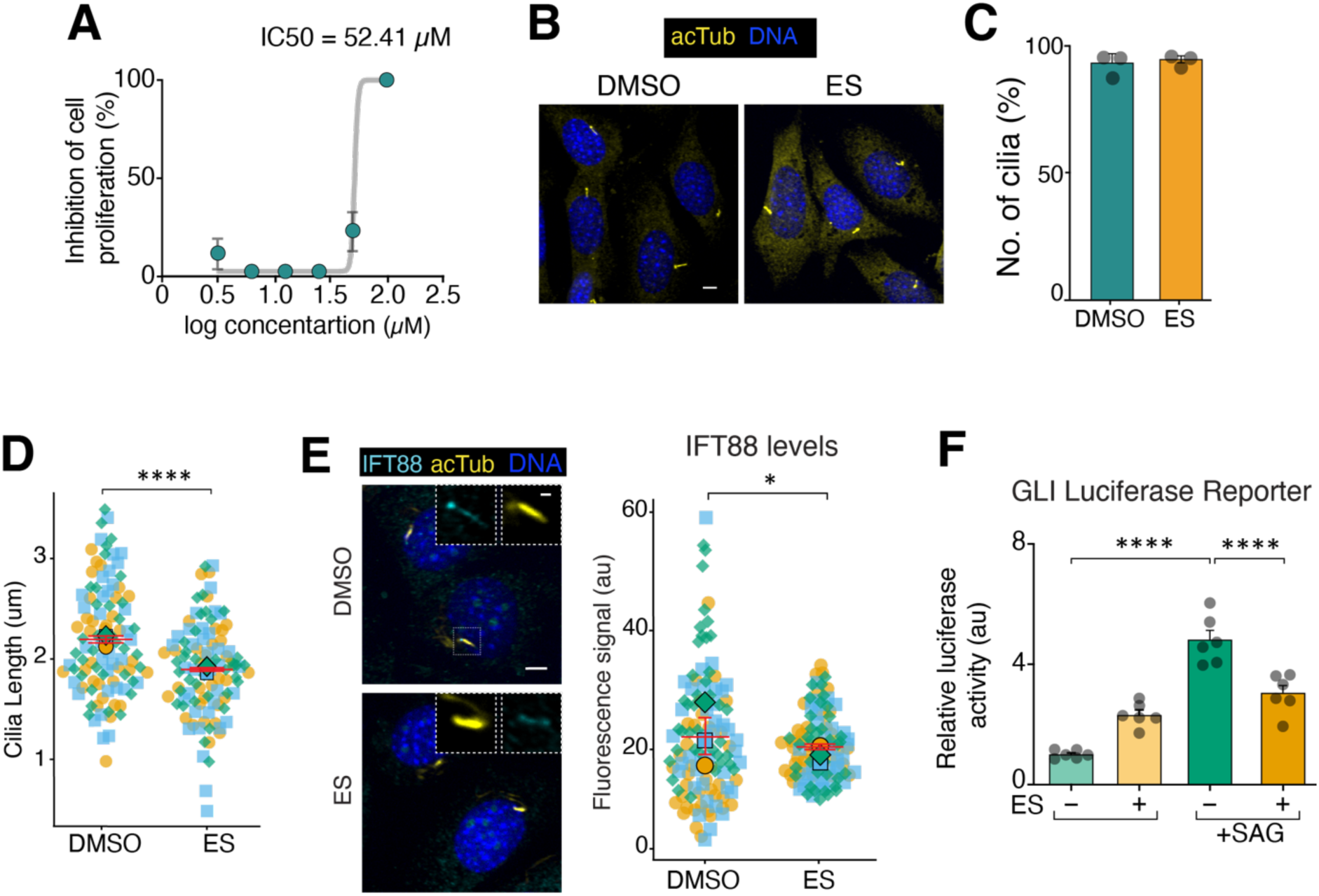
Effect of endosulfan on cilia and *GLI* expression in NIH3T3. **(A)** Cytotoxicity assay showing IC50 determination for endosulfan in NIH3T3 cells after 48 h of endosulfan treatment. **(B)** NIH3T3 cells treated as indicated were stained for acTub (yellow) and DNA (blue). **(C)** Percentage of ciliated cells quantified from (**B**). Data represent the percentage of ciliated cells from three independent experiments, mean ± SD. **(D)** Quantification of ciliary length measured from acTub (panel 2). Superplots represent all data points from three independent experiments ±SEM. Cell numbers: DMSO, n = 102; endosulfan, n = 103. **(E)** Representative immunofluorescence micrographs showing IFT88 (cyan), acTub (yellow) and DNA (blue). Quantification of ciliary IFT88 levels (panel 2) in NIH3T3 cells treated with endosulfan or vehicle following serum starvation. Cell numbers: DMSO n = 115, endosulfan n = 109. **(F)** GLI-luciferase reporter activity in NIH3T3-Shh-LightII cells treated with endosulfan or vehicle. Data represent the mean ±SD of six independent experiments. Fluorescence intensity is expressed in arbitrary units (a.u.). Scale bars, 5 µm; inset scale bars, 1 µm. Asterisks denote statistical significance as determined by unpaired Student’s t-test or one-way ANOVA followed by Tukey’s post-hoc multiple comparison test, as appropriate: **** p ≤ 0.0001; *** p ≤ 0.001; ** p ≤ 0.01; * p ≤ 0.05.

**Figure EV4:**
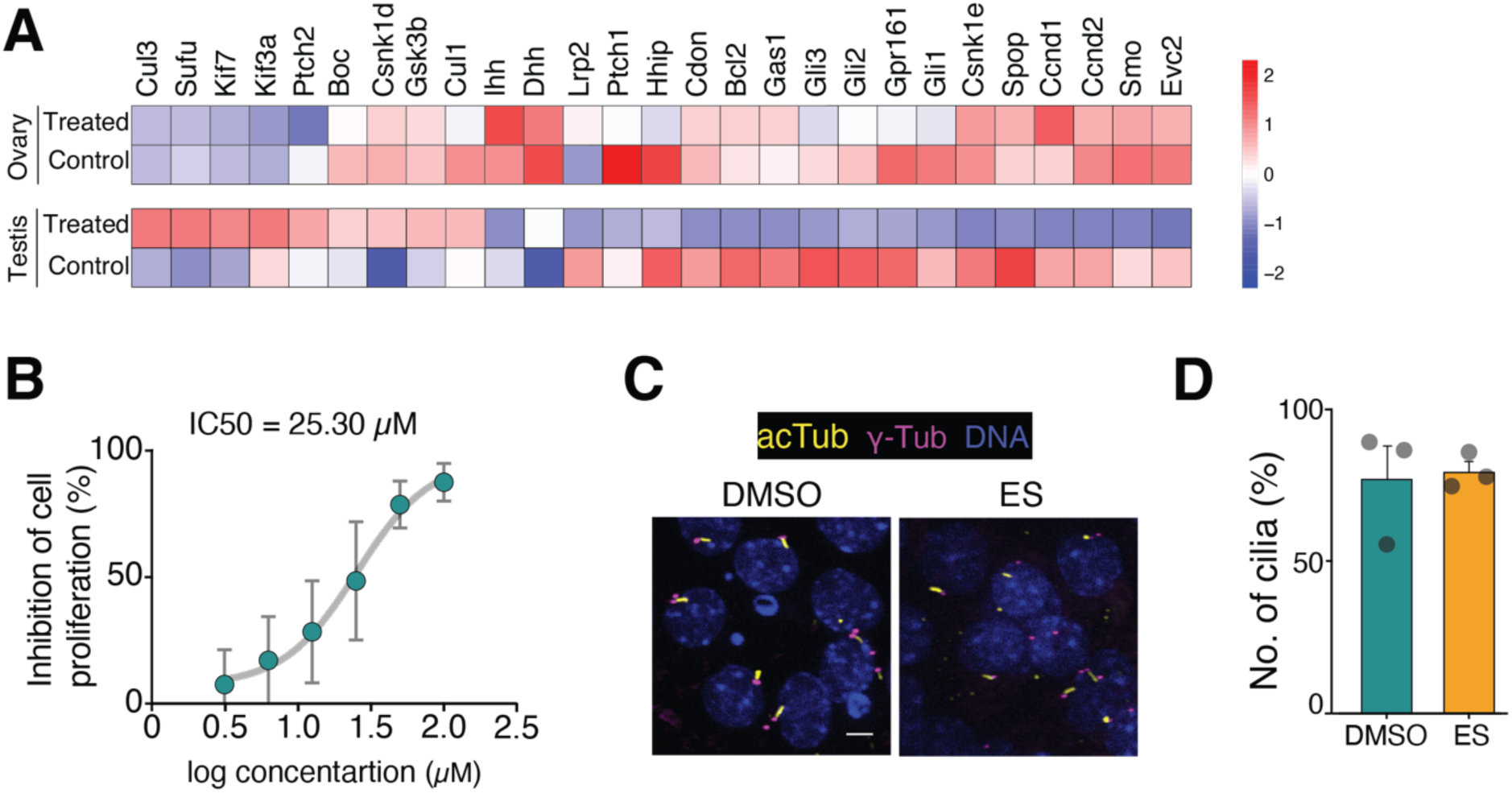
Effect of endosulfan on cilia and Hedgehog pathway genes in gonads. **(A)** Heatmap showing RNA-seq expression data for hedgehog pathway genes from the gonads of endosulfan-exposed mice. Expression values were log2-transformed [log2(X + 1)]; genes exhibiting zero variance across all samples were excluded prior to visualisation. Colour intensity represents the Z-score of normalised expression values. **(B)** Cytotoxicity assay showing IC_50_ determination for endosulfan in TM3 cells following 48 h endosulfan treatment. **(C and D)** Representative immunofluorescence micrographs showing acetylated tubulin (acTub, yellow), γ-tubulin (γ-Tub, magenta), and DNA (blue) **(C)** and quantification of the percentage of ciliated cells **(D)** in TM3 cells treated with endosulfan or vehicle. Data represent the percentage of ciliated cells from three independent experiments ±SD.

**Figure EV5:**
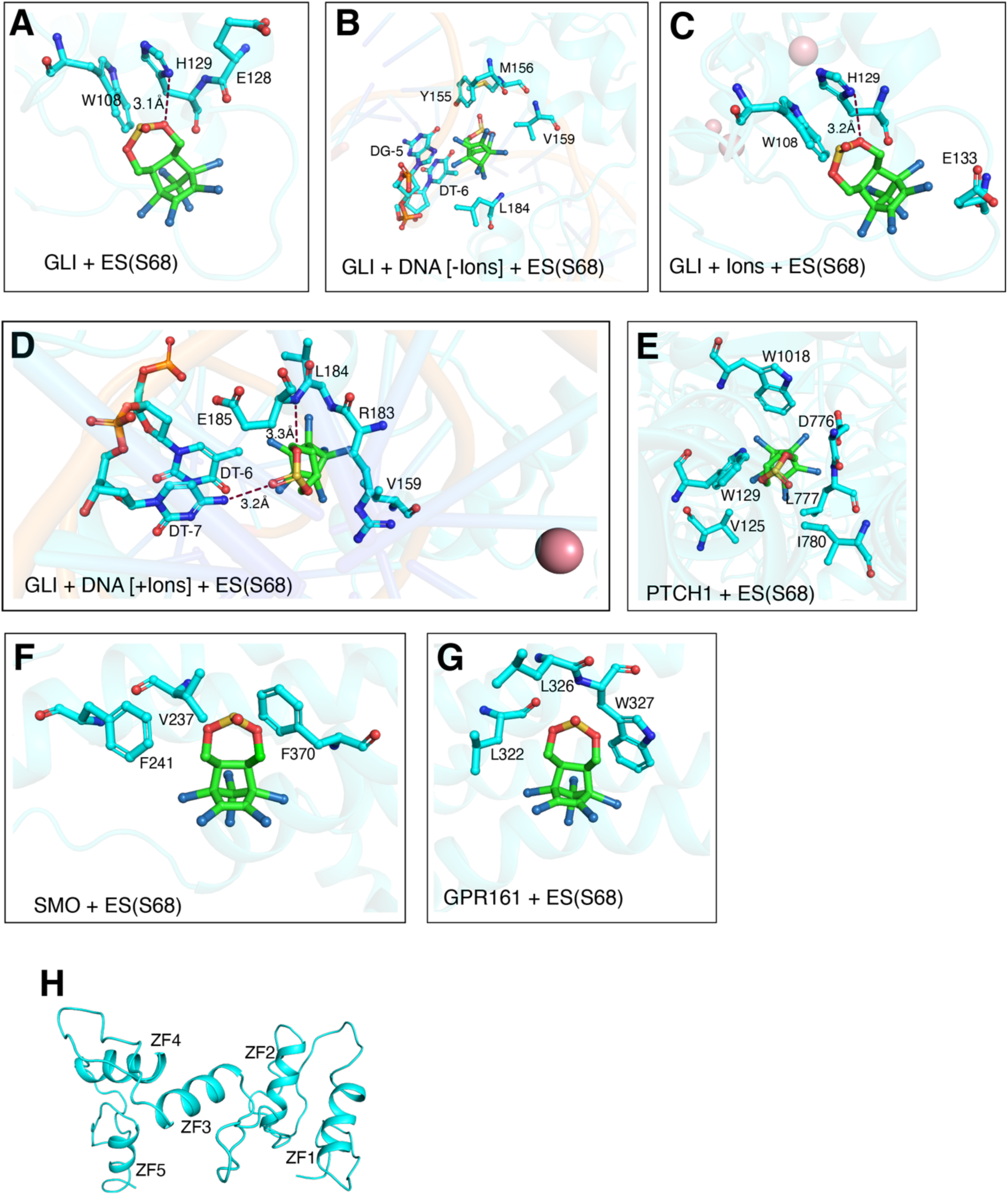
Endosulfan binding with Hedgehog pathway components. **(A-G)** Representative docking poses of endosulfan at the binding sites of GLI without DNA and ions (**A**), GLI without ions (**B**), GLI without DNA (**C**), GLI–DNA complex (**D**), PTCH1 (**E**), SMO (**F**), and GPR161 (**G**). Key interacting residues are labelled; hydrogen bonds are represented as dashed lines in (Å). **(H)** Representative pose of GLI.

**Figure EV6:**
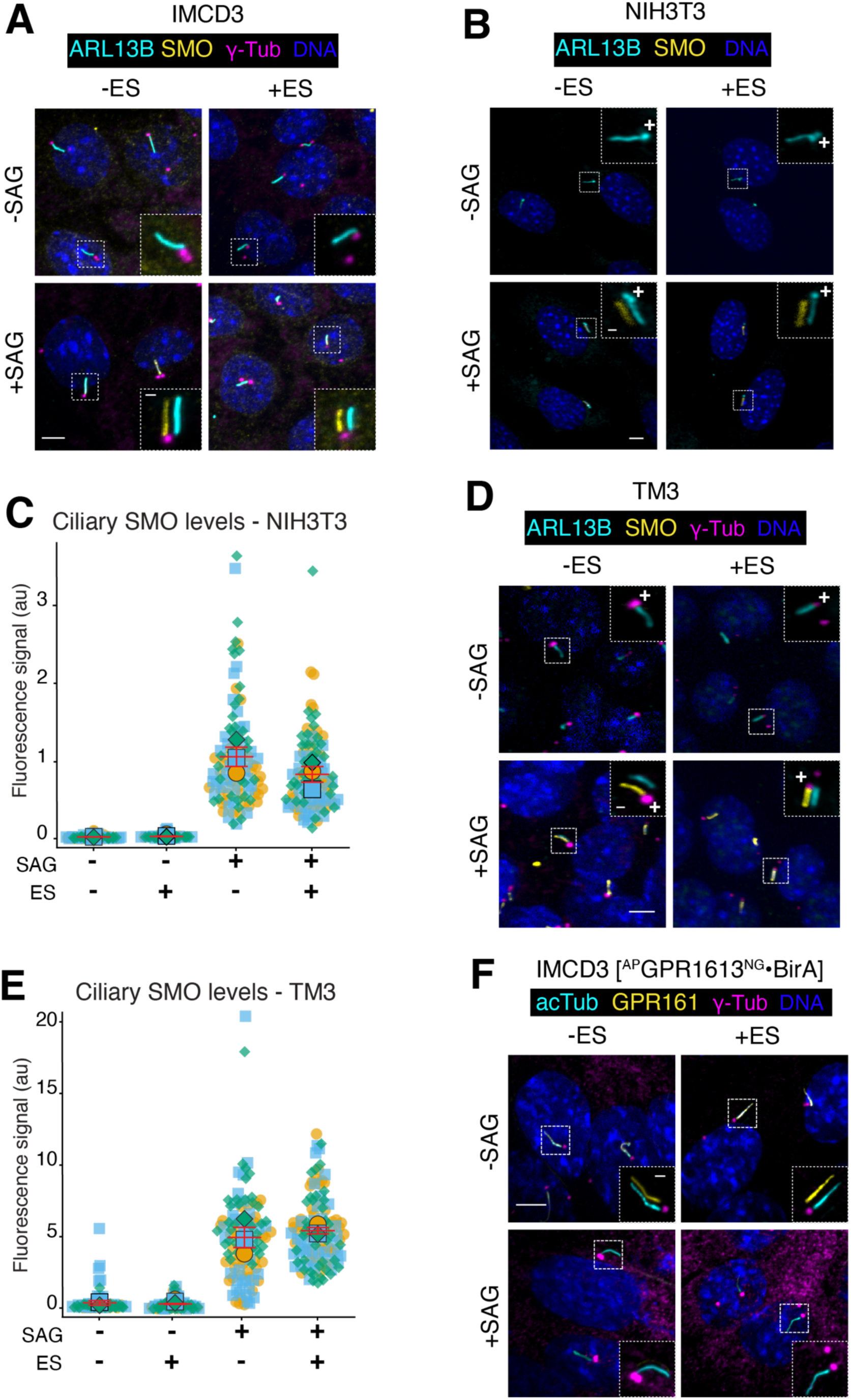
Effect of endosulfan on SMO and GPR161 ciliary localisation. **(A)** Representative immunofluorescence micrographs corresponding to Figure 3C. **(B and C)** Representative immunofluorescence micrographs showing ARL13B (cyan) and SMO (yellow) **(B)** and quantification of ciliary SMO levels **(C)** in NIH3T3 cells treated with endosulfan or vehicle in the presence or absence of SAG. Data represent all data points from three independent experiments ±SEM. Cell numbers: DMSO n = 110, endosulfan n = 109, SAG n = 116, SAG + endosulfan n = 109. **(D and E)** Representative immunofluorescence micrographs showing ARL13B (cyan), SMO (yellow), and γ-Tub (magenta) **(D)** and quantification of ciliary SMO levels **(E)** in TM3 cells treated with endosulfan or vehicle in the presence or absence of SAG. Superplot convention as in **(C)**. Data represent all data points from three independent experiments ±SEM. Cell numbers: DMSO n = 102, endosulfan n = 104, SAG n = 109, SAG + endosulfan n = 120. **(F)** Representative immunofluorescence micrographs showing acetylated tubulin (acTub, cyan), GPR161 (yellow), and γ-Tub (magenta) in IMCD3-FlpIn cells treated as indicated. Fluorescence intensity is expressed in arbitrary units (a.u.). Scale bars, 5 µm; inset scale bars, 1 µm. Asterisks denote statistical significance as determined by unpaired Student’s t-test or one-way ANOVA followed by Tukey’s post-hoc multiple comparison test, as appropriate: **** p ≤ 0.0001; *** p ≤ 0.001; ** p ≤ 0.01; * p ≤ 0.05.

**Figure EV7:**
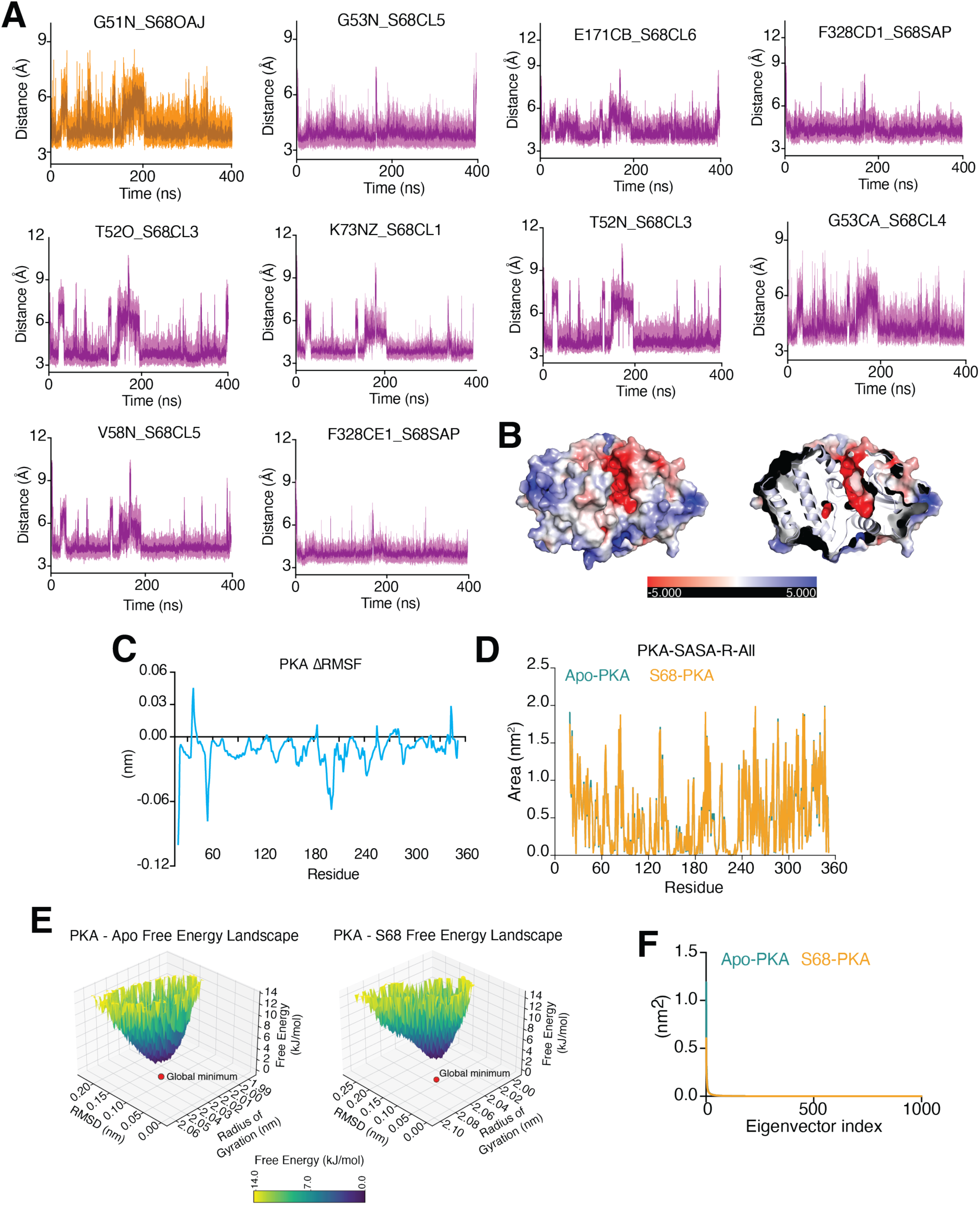
Endosulfan interacts stably with PKA-C throughout the MD simulation. **(A)** Distance–time plots of key PKA-C residues interacting with endosulfan over the 400 ns simulation. Orange traces represent hydrogen-bond distances; purple traces represent hydrophobic interaction distances (Å). The full MD simulation trajectory is provided in Movie **S3**. **(B)** Electrostatic surface potential representation of the PKA-C apo state. **(C)** Delta RMSF (ΔRMSF) for individual PKA-C residues (nm), illustrating changes in residue fluctuations. **(D)** Total solvent-accessible surface area (SASA) of apo (green) and endosulfan-bound (wheat) PKA-C over the course of the simulation, reflecting the overall change in solvent accessibility upon endosulfan binding. **(E)** Three-dimensional free energy landscape (3D-FEL) of PKA-C in the apo and endosulfan-bound states, illustrating the conformational energy states sampled during the simulation. The global energy minimum of each state is indicated by a red dot. **(F)** Conformational entropy analysis of PKA-C in the apo and endosulfan-bound states, showing that endosulfan binding does not alter global conformational entropy.

**Figure EV8:**
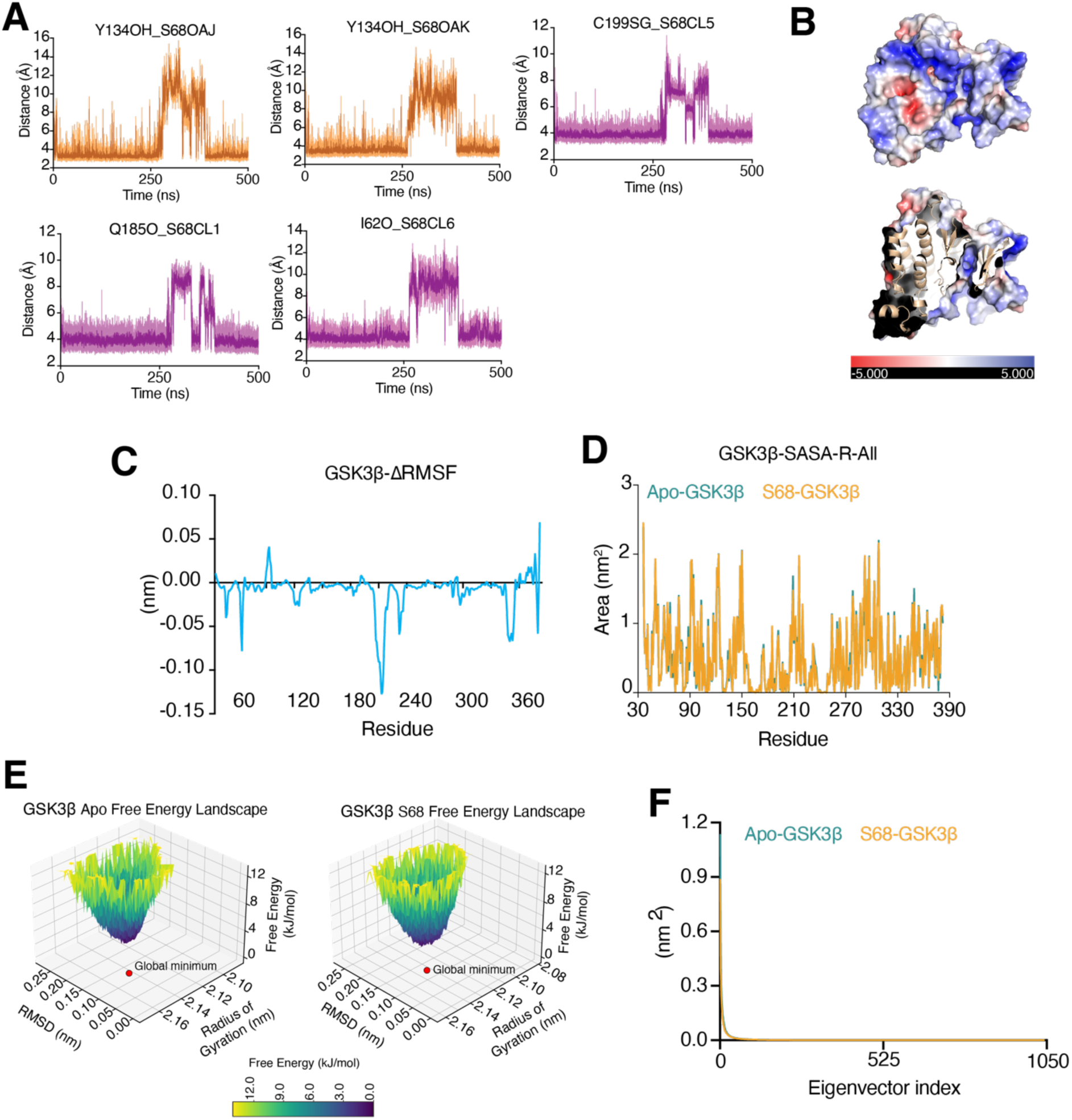
Endosulfan interacts stably with GSK3β throughout the MD simulation. **(A)** Distance–time plots of key GSK3β residues interacting with endosulfan over the 400 ns simulation. Orange traces represent hydrogen-bond distances; purple traces represent hydrophobic interaction distances (Å). The full MD simulation trajectory is provided in Movie **S4**. **(B)** Electrostatic surface potential representation of the GSK3β apo state. **(C)** Delta RMSF (ΔRMSF) for individual GSK3β residues (nm), illustrating changes in residue fluctuations. **(D)** Total solvent-accessible surface area (SASA) of apo (green) and endosulfan-bound (wheat) GSK3β over the course of the simulation, reflecting the overall change in solvent accessibility upon endosulfan binding. **(E)** Three-dimensional free energy landscape (3D-FEL) of GSK3β in the apo and endosulfan-bound states, illustrating the conformational energy states sampled during the simulation. The global energy minimum of each state is indicated by a red dot. **(F)** Conformational entropy analysis of GSK3β in the apo and endosulfan-bound states, showing that endosulfan binding does not alter global conformational entropy.

**Figure EV9:**
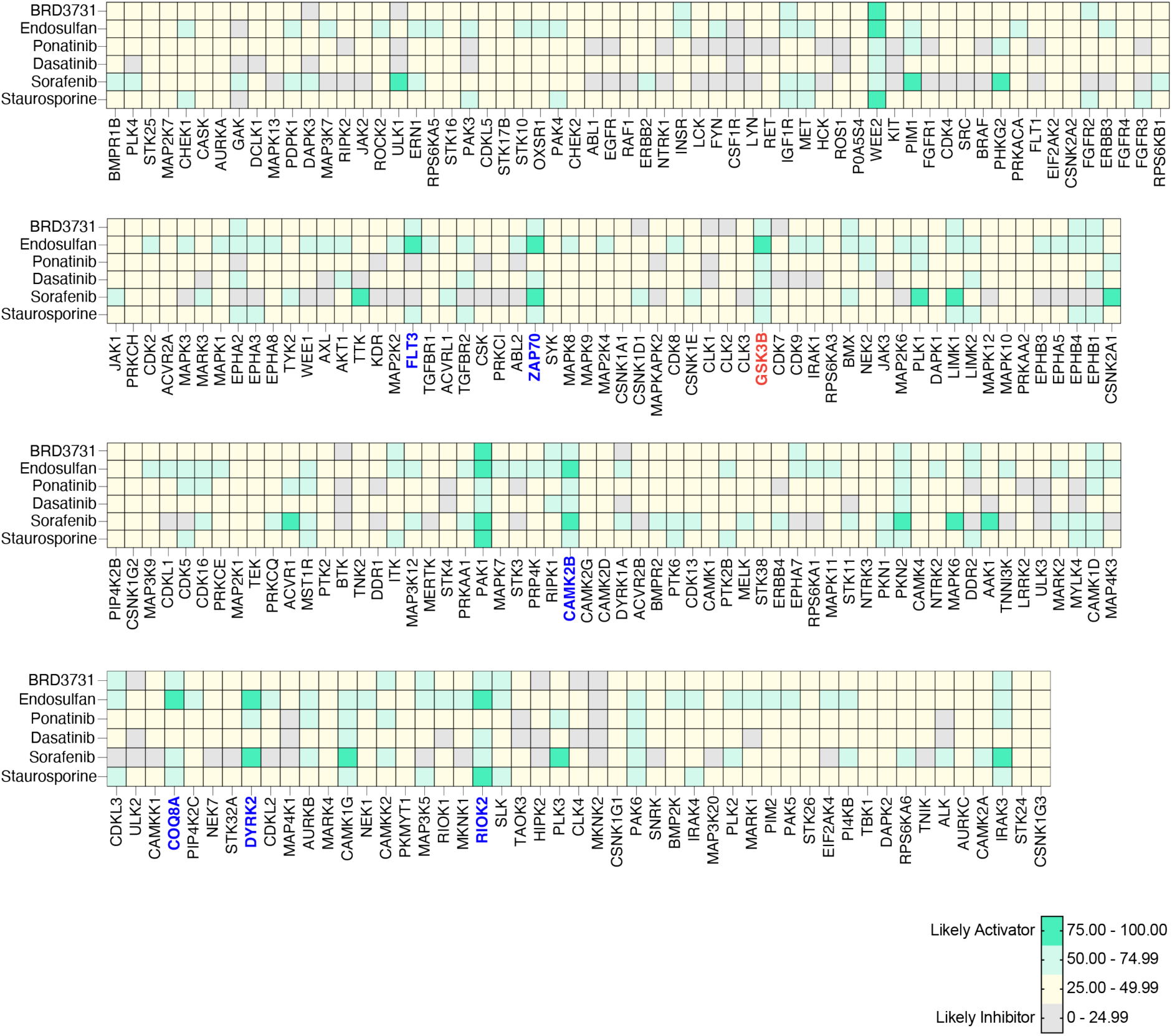
Binding affinity of endosulfan to kinases compared with kinase inhibitors. A binding affinity heatmap generated by DeepKinomeWeb shows predicted compound–kinase binding affinities across a predefined kinase panel. Each cell represents a compound–kinase pair; darker green indicates lower predicted percentage inhibition, reflecting higher binding affinity. Endosulfan showed binding affinity similar to known kinase inhibitors for the kinases shown in blue.

**Figure EV10:**
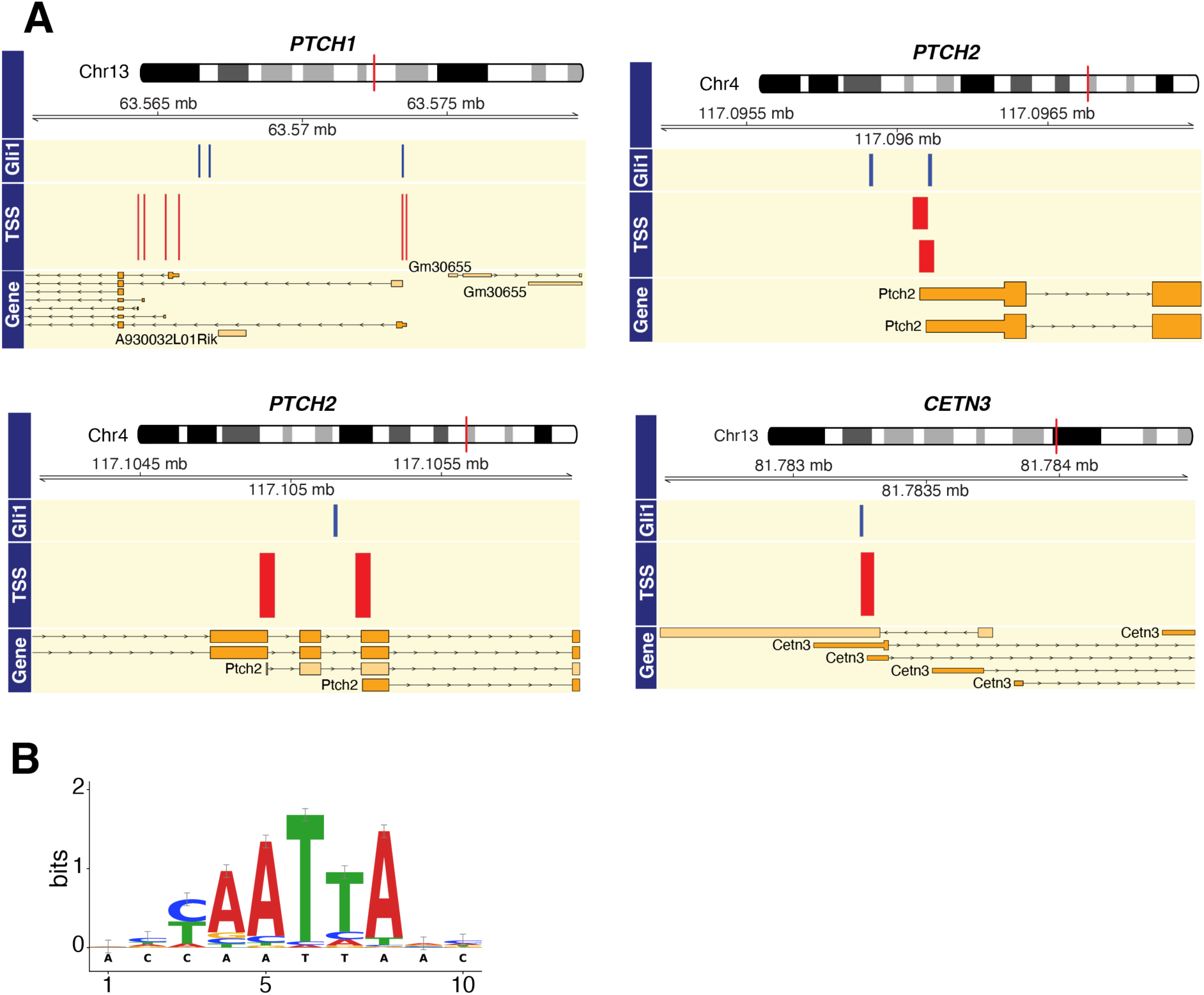
GLI1 binding motif analysis from ChIP-seq and MEME Suite. **(A)** Genome browser views of ChIP-seq signal at Hedgehog pathway and ciliary gene loci. Tracks show the ChIP-seq signal alongside TSS-associated peaks and gene annotations at the following loci: Ptch1 on chromosome 13 (∼63.57 Mb), Ptch2 on chromosome 4 (∼117.09 Mb and ∼117.10 Mb), and Cetn3 on chromosome 13 (∼81.78 Mb). The red vertical line marks the gene position within each locus. **(B)** Consensus sequence logo of the GLI1 binding motif derived from MEME Suite analysis of candidate target gene promoters. Letter height reflects information content (bits) at each nucleotide position, indicating the degree of conservation; error bars above each letter represent positional uncertainty estimated from multiple aligned sequences.

**Figure EV11:**
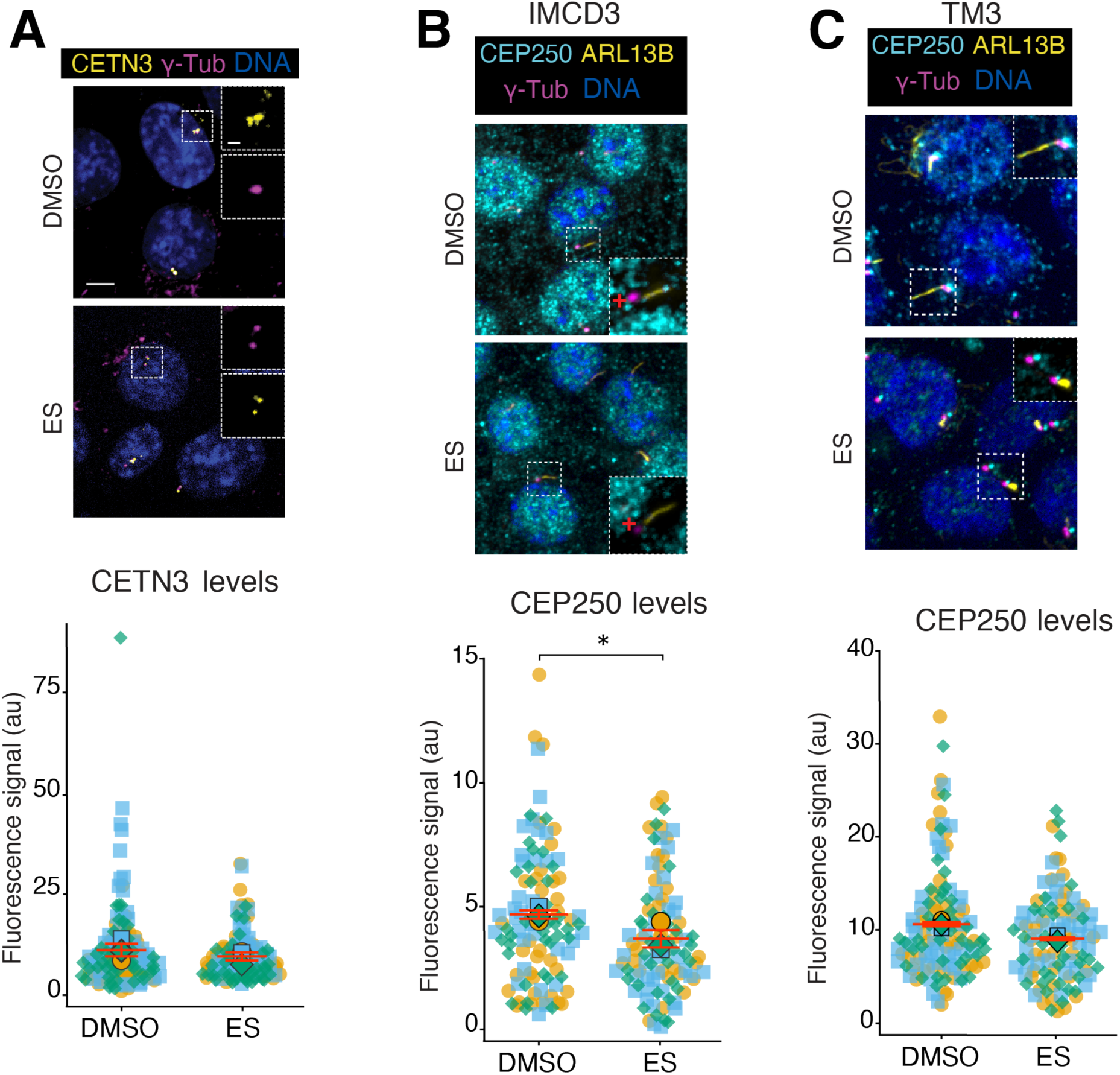
Effect of endosulfan on Centrin3 and CEP250. **(A)** Representative immunofluorescence micrographs showing Centrin3 (yellow) and γ-tubulin (γ-Tub, magenta), and quantification of CETN3 levels at the ciliary base in IMCD3 cells treated with endosulfan or vehicle. In the superplots, colours indicate individual experiments; each dot represents a single cell, and the larger dot indicates the experiment mean. Data represent all data points from three independent experiments ±SEM. Cell numbers: DMSO n = 140, endosulfan n = 130. **(B)** Representative immunofluorescence micrographs showing CEP250 (Cyan), ARL13B (yellow), and γ-tubulin (γ-Tub, magenta), and quantification of CEP250 levels at the ciliary base in IMCD3 cells treated with endosulfan or vehicle. Superplot convention as in **A**. Data represent all data points from three independent experiments ±SEM. Cell numbers: DMSO n = 106, endosulfan n = 101. **(C)** Representative immunofluorescence micrographs showing CEP250 (Cyan), ARL13B (yellow), and γ-tubulin (γ-Tub, magenta), and quantification of CEP250 levels at the ciliary base in TM3 cells treated with endosulfan or vehicle. Superplot convention as in **A.** Data represent all data points from three independent experiments ±SEM. Cell numbers: DMSO n = 120, endosulfan n = 106. Fluorescence intensity is expressed in arbitrary units (a.u.). Scale bars, 5 µm; inset scale bars, 1 µm. Asterisks denote statistical significance as determined by unpaired Student’s t-test or one-way ANOVA followed by Tukey’s post-hoc multiple comparison test, as appropriate: **** p ≤ 0.0001; *** p ≤ 0.001; ** p ≤ 0.01; * p ≤ 0.05.

**Table EV1:** Ciliopathy phenotypes in Zebrafish.

| Model System | Phenotypes / Effects | Reference (PMID) |
| --- | --- | --- |
| Zebrafish (Embryo) | Dose-dependent embryonic deformities including edema malformations, pericardial edema, reduced pigmentation; larval hyperactivity (developmental neurotoxicity) at 6 dpf | PMID: 28065758 - Pereira et al., 2017 |
| Zebrafish (Embryo) | Morphological deformities (yolk/pericardial edema, spinal curvature, tail defects); seizure events; abnormal midbrain/hindbrain development; inhibited heart development and reduced cardiac function | PMID: 39929004 - Kin et al., 2025 |
| Zebrafish (Embryo/Larva) | Developmental neurotoxicity; abnormal touch response; inhibition of touch-response at low waterborne concentrations | PMID: 19883949 - Herman et al., 2010 |
| Zebrafish (Embryo) | Arrested epiboly progress, shortened body length, curved trunk, incomplete/malformed brain structure; effects enhanced at elevated temperature (35°C) | PMID: 37188935 - Atta et al., 2023 |
| Zebrafish (Embryo/Adult) | Endocrine disruption; decreased hatching rate, testicular pathology in males, reduced gonadosomatic index in females, elevated vitellogenin in males | PMID: 21726828 - Han et al., 2011 |
| Zebrafish (Adult) | Inhibition of brain AChE activity; impaired swimming performance; reduced exploration; impaired long-term memory formation | PMID: 22450085 - Pereira et al., 2012 |
| Zebrafish (Adult) | DNA damage (comet assay); oxidative stress; depletion of antioxidant enzyme capacity (CAT, SOD); ROS overproduction | PMID: 22535316 - Shao et al., 2012 |
| Zebrafish (Adult + Offspring) | Hepatocyte enlargement; reduced red blood cell counts; increased oxidative stress (TBARS); reduced offspring heart rate and altered morphometrics | PMID: 21262189 - Valencia-Santamaria et al., 2011 |
| Zebrafish (Adult) | Induction of hepatic CYP enzymes and glutathione-S-transferases; sex-based differences in enzyme induction | PMID: 23523001 - Han et al., 2003 |
| Zebrafish<br>(Embryo) | Endosulfan sulfate metabolite causes spinal curvature, seizure-like activity, pericardial edema, heart-development inhibition | PMID: 34455181 -<br>Lee et al., 2021 |

**Table EV2:** Ciliopathy phenotypes in Mouse.

| Model System | Phenotypes / Effects | Reference (PMID) |
| --- | --- | --- |
| Mouse (Male) | Testicular atrophy; impaired spermatogenesis; increased ROS in epididymis; reduced sperm count and motility; male-specific infertility | PMID: 27551436<br>- Sebastian & Rajasekaran, 2016 |
| Mouse (Male & Female) | Long-term reproductive toxicity; testicular vacuole degeneration; loss of Graafian follicles; DNA damage persisting months after exposure; tumor formation | PMID: 36506329<br>- Sharma et al., 2022 |
| Mouse (Male & Female, Swiss Albino) | Degeneration of germinal epithelium, Graafian follicles and corpus luteum; seminiferous tubule damage; altered hormonal profiles; chromosomal aberrations | PMID: 38783074<br>- Kumar et al., 2024 |
| Mouse (Male, Developmental Exposure) | Nigrostriatal dopaminergic damage; reduction in dopamine transporter and tyrosine hydroxylase; increased susceptibility to MPTP challenge (Parkinson's disease model) | PMID: 25002410<br>- Gaur et al., 2014 |
| Mouse (Male, Reproductive) | Mitochondrial dysfunction in spermatogenic cells; sperm tail breakage; reduced sperm count and quality via oxidative-stress mechanisms | PMID: 22705298<br>- Wang et al., 2012 |
| Mouse (C57BL/6) | Hepatotoxicity; hepatocyte ROS accumulation; stellate-cell activation; immune activation/inflammation; disrupted hepatic microenvironment | PMID: 40442041<br>- Bai et al., 2025 |
| Mouse (Swiss Albino) | Hepatic neoplastic changes; vacuolation of sinusoidal spaces; hemorrhage; Kupffer-cell proliferation; ultrastructural degeneration in hepatocytes | SCIRP: 47613 - Siddiqui et al., 2014 |

**Table EV3:**
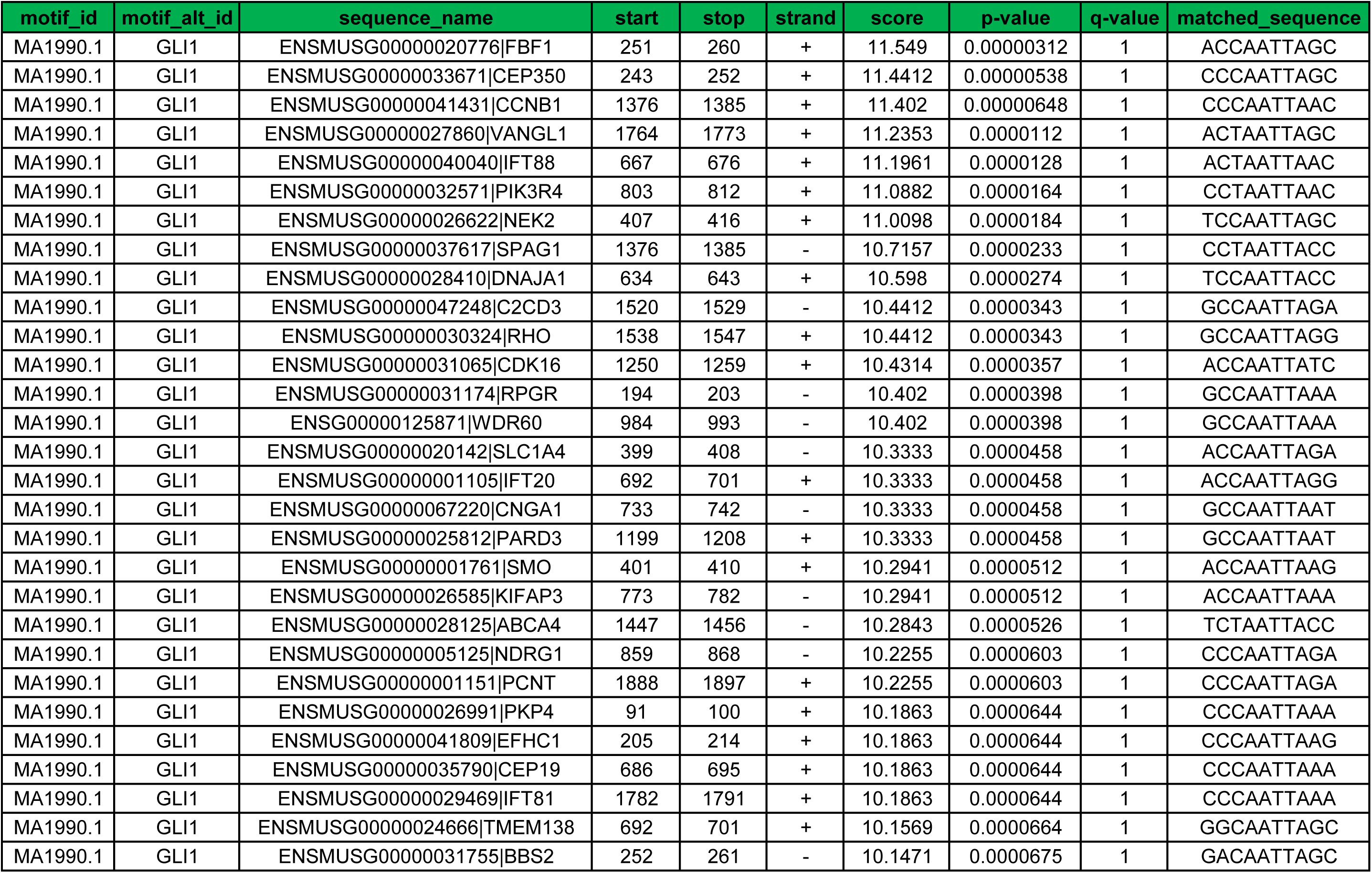
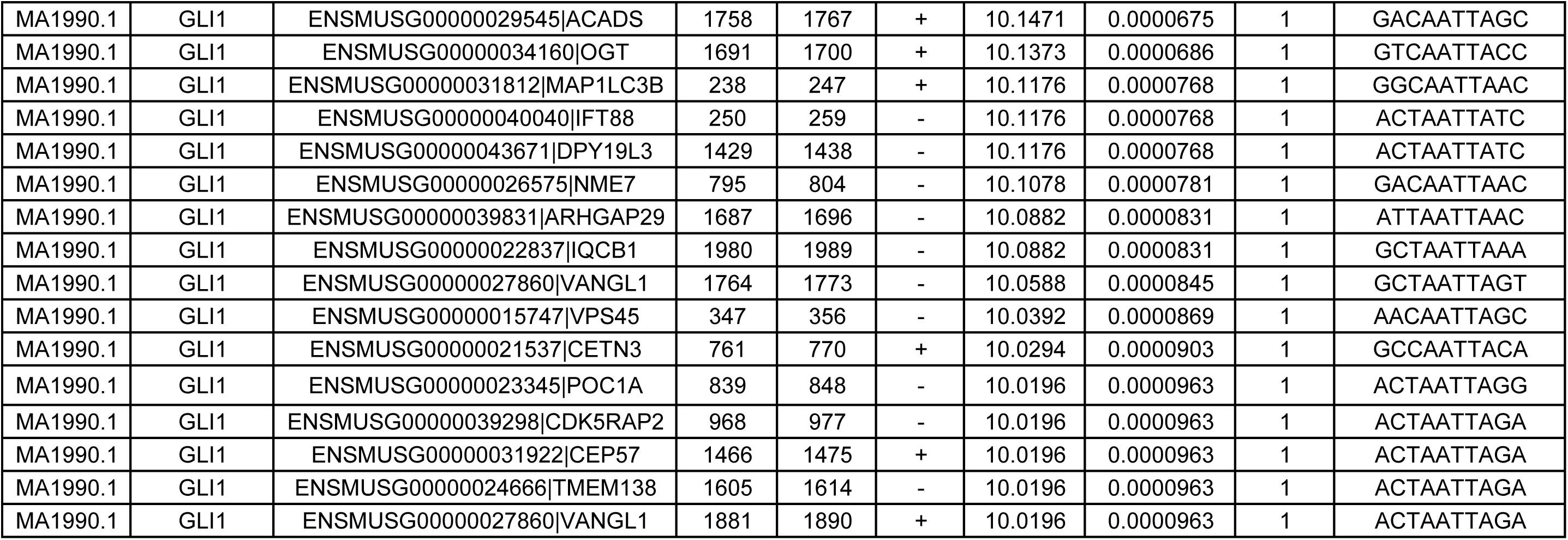

**Table EV4:**
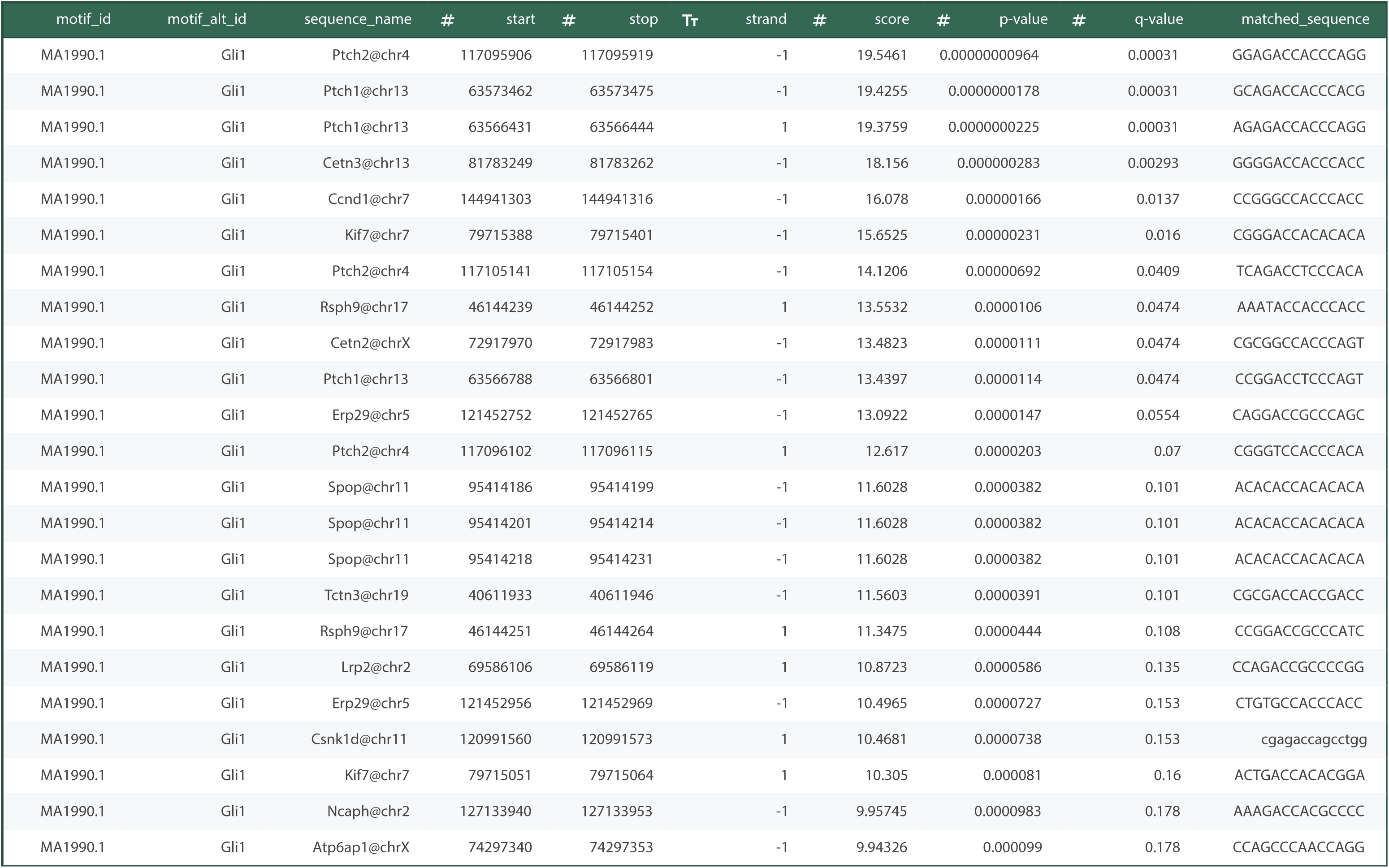

